# The WW domain of IQGAP1 binds directly to the p110α catalytic subunit of PI 3-kinase

**DOI:** 10.1101/2022.09.24.509339

**Authors:** A. Jane Bardwell, Madhuri Paul, Kiku C. Yoneda, Maria D. Andrade-Ludeña, Oanh T. Nguyen, David A. Fruman, Lee Bardwell

**Affiliations:** Dept. of Developmental and Cell Biology, University of California, Irvine, CA, USA; Dept. of Molecular Biology and Biochemistry, University of California, Irvine, CA, USA

**Keywords:** Scaffold, PI3K, Phosphoinositide, Signaling, Cancer

## Abstract

IQGAP1 is a multi-domain cancer-associated protein that serves as a scaffold protein for multiple signaling pathways. Numerous binding partners have been found for the calponin homology, IQ and GAP-related domains in IQGAP1. Identification of a binding partner for its WW domain has proven elusive, however, even though a cell-penetrating peptide derived from this domain has marked anti-tumor activity. Here, using *in vitro* binding assays with human proteins and co-precipitation from human cells, we show that the WW domain of human IQGAP1 binds directly to the p110α catalytic subunit of phosphoinositide 3-kinase (PI3K). In contrast, the WW domain does not bind to ERK1/2, MEK1/2, or the p85α regulatory subunit of PI3K when p85α is expressed alone. However, the WW domain is able to bind to the p110α/p85α heterodimer when both subunits are co-expressed, as well as to the mutationally activated p110α/p65α heterodimer. We present a model of the structure of the IQGAP1 WW domain, and experimentally identify key residues in the hydrophobic core and beta strands of the WW domain that are required for binding to p110α. These findings contribute to a more precise understanding of IQGAP1-mediated scaffolding, and of how IQGAP1-derived therapeutic peptides might inhibit tumorigenesis.

## Introduction

Recently there has been renewed enthusiasm for targeting protein-protein interactions as a therapeutic strategy for treating cancer and other diseases, and for using peptide-based inhibitors as drugs or drug leads in such approaches [1–11]. Signaling scaffold proteins – large, multidomain proteins that bind to multiple members of one or more signaling pathways – participate in numerous protein-protein interactions that could be targeted to disrupt signaling [7, 12–16]. Indeed, studies with human IQGAP1, a scaffold protein for multiple cancer-associated signaling pathways [17–21], have led to the discovery that its 26-residue WW domain, when engineered as a cell-penetrating peptide, shows broad anti-tumor activity with minimal associated toxicity [12, 22].

WW domains are one of the smallest independently folding protein interaction domains, and thus have a strong potential as leads in peptidomimetic development. These domains are typically between 25 and 35 residues in length and fold into a three-stranded antiparallel beta-sheet [23–25]. WW domains are characterized as proline recognition domains, and typically bind to proline-rich short linear motifs in their binding partners [26]. A cell-penetrating derivative of the IQGAP1 WW domain was first developed by Jameson *et al*. [12], who showed that this peptide inhibited the proliferation, migration, viability and tumor-forming ability of breast, colorectal and melanoma tumor cells that contained activating mutations in the RAS/MAPK pathway. Non-transformed or primary cells were unaffected. They further showed that the WW peptide, when delivered systemically, significantly increased the lifespan of mice bearing pancreatic tumors [12]. Choi *et al*. [22] showed that the WW peptide reduced the viability of multiple additional breast cancer cell lines, but did not adversely affect normal control lines. The anti-tumor activity of the WW peptide has also been confirmed in other studies [27, 28].

In view of these exciting results, it is imperative to determine the binding partner(s) of the WW domain. For many years, it was believed that the ERK1 and ERK2 MAP kinases bound to the WW domain of IQGAP1, and were the only proteins to do so [29–31]. Indeed, this belief motivated the Jameson *et al*. study, as these workers sought to target ERK signaling by using the WW domain as a competitive inhibitor of ERK-IQGAP1 binding. In 2017, however, we showed that ERK1 and ERK2 do not bind to IQGAP1’s WW domain, but instead bind to the nearby IQ domain [32]. This finding revived the search for the authentic target of the IQGAP1 WW domain.

A significant clue in this regard came from a landmark 2016 study from the Anderson lab [22]. These authors showed that IQGAP1 acts a scaffold protein for the phosphoinositide 3-kinase (PI3K) pathway by binding to several enzymes that modify phosphoinositides, including PI3K itself. PI3Ks are a family of enzymes that phosphorylate inositol lipids in response to signals from growth factors, cytokines, and other cellular and environmental cues. In so doing, they regulate cell proliferation, growth, survival, motility, intracellular trafficking, and metabolism [33–37]. The most common PI3K isoform is a heterodimer of a catalytic subunit designated p110α (gene *PIK3CA*) and a regulatory subunit designated p85α (gene *PIK3R1*). Following activation by growth factors, insulin, or other signals, the p110α/p85α holoenzyme phosphorylates the inositol ring of phosphatidylinositol-(4,5)-biphosphate (PIP2) on the 3-position to generate phosphatidylinositol-(3,4,5)-triphosphate (PIP3), a key intracellular second messenger. Both subunits of PI3K, particularly p110α, are recognized oncogenic drivers that are frequently mutated in human cancer and tissue overgrowth syndromes. In addition to being activated by direct mutation, PI3K activity can also be pathologically activated by other means, including mutations in other oncogenes, loss of tumor suppressor genes (especially the PTEN tumor suppressor), and oncovirus-mediated transformation [33–37]. Hence, PI3K activation is a hallmark of cancer [35].

Here, we show that the WW domain of human IGQAP1 binds directly to human p110α protein, the catalytic subunit of the PI3K holoenzyme, but does not bind to p85α, ERK1, ERK2, MEK or MEK2. We also show that the anti-tumor WW peptide has these same binding specificities. In addition, we present a model of this WW domain, and investigate which residues in the domain participate in the interaction with p110α. These findings contribute to a more precise understanding of IQGAP1-mediated scaffolding, and of how IQGAP1-derived therapeutic peptides might inhibit tumorigenesis.

## Results

### The WW domain of IQGAP1 binds specifically to p110α

Although IQGAP1 had been shown to act as a scaffold protein for the PI 3-kinase pathway [22] and the RAS/RAF/MEK/ERK pathway [29, 32], the precise binding partner of IQGAP1’s WW domain was unknown. We took a “candidate protein” approach to finding this partner, using components from both of these pathways. First, the WW domain of human IQGAP1 (residues 684-710, Figure 1A) was fused at its N terminus to *Schistosoma japonicum* glutathione *S*-transferase (GST), and the resulting fusion protein (GST-WW) was expressed in bacteria and purified by adsorption to glutathione-Sepharose beads. Next, GST-WW (or GST alone as a negative control) was incubated with full-length human p85α or p110α that had been produced in radiolabeled form by *in vitro* translation (Figure 1B). We also assessed the ability of GST-WW to bind to *in vitro* translated human ERK2 protein (Figure 1B). Bead-bound complexes were collected by sedimentation, washed extensively, and analyzed by SDS-PAGE and autoradiography. As shown in Figure 1C, full-length p110α bound efficiently to WW, but p85α and ERK2 did not. Furthermore, the binding of p110α was specific, because only trace precipitation of p110α occurred when GST was used instead of the GST-WW fusion protein. As an additional control, we tested two other small GST fusion protein fragments for binding to p110α, neither of which bound above background (Supplementary Figure 1).

**Figure 1.**
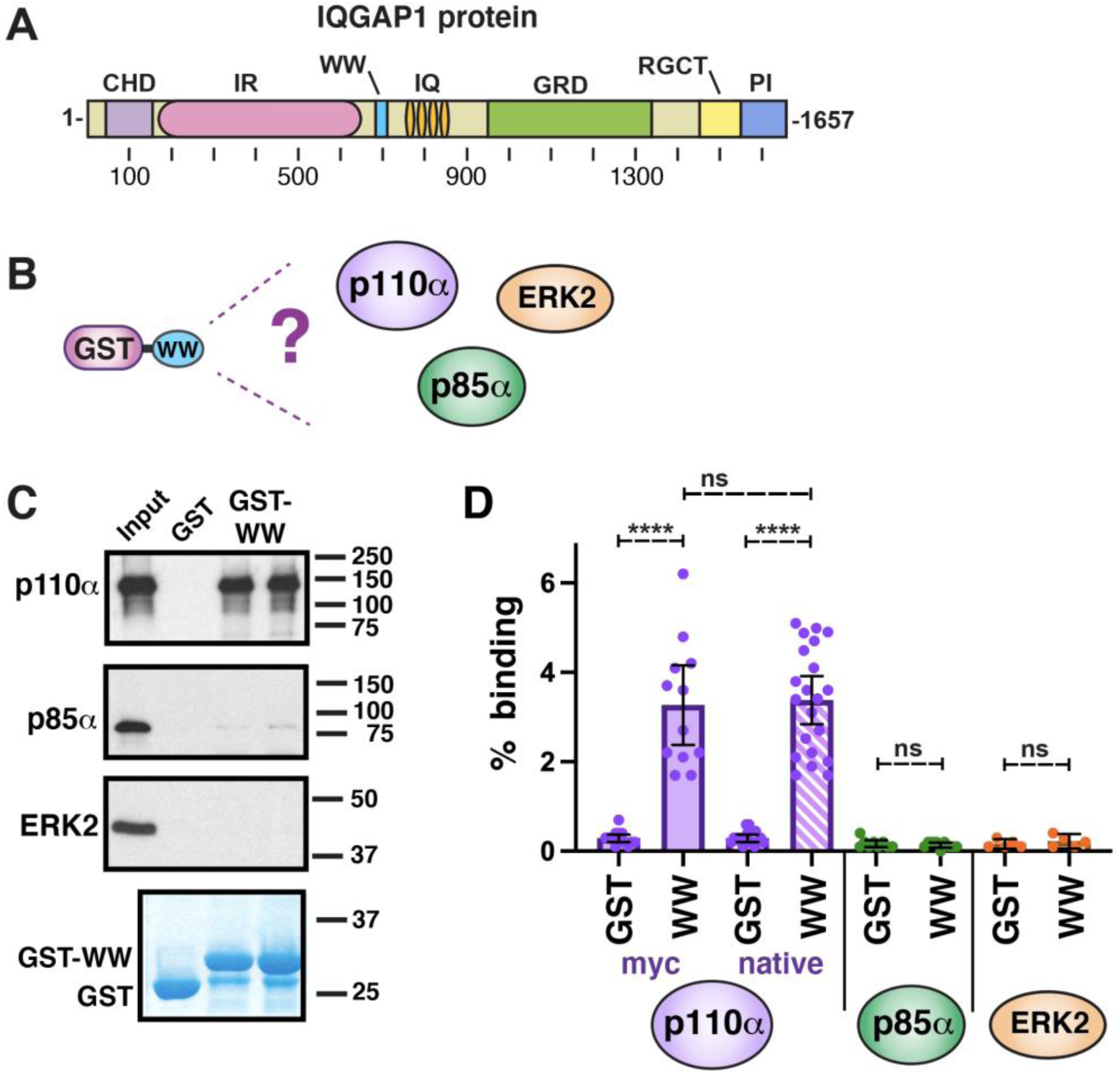
The WW domain of IQGAP binds to the catalytic subunit of PI 3-kinase. (**A**) Schematic depicting full-length human IQGAP1 protein with domains denoted as follows: CHD, calponin homology domain; IR, internal repeated sequence/coiled-coil domain; WW, WW domain; IQ, IQ domain; GRD, GTPase-activating protein-related domain; RGCT, RasGAP C-terminal domain; PI, phosphoinositide binding domain. (**B**) The WW domain of human IQGAP1, fused to GST, was tested for binding to the p110α and p85α subunits of PI3K, as well as to the ERK2 MAP kinase. (**C**) The WW domain binds to p110α, but not to p85α or ERK2. As shown in **B**, ^35^S-radiolabeled fulllength human p110α, p85α and ERK2 proteins were prepared by *in vitro* translation and partially purified by ammonium sulfate precipitation, and portions (5% of the amount added in the binding reactions) were resolved on a 10% SDS-polyacrylamide (SDS-PAGE) gel; these samples are labeled “Input” in the figure. Portions (1 pmol) of the same proteins were incubated with 25 μg of GST or GST-WW bound to glutathione-Sepharose beads, and the resulting bead-bound protein complexes were isolated by sedimentation and resolved by 10% SDS-PAGE on the same gel. The gel was analyzed by staining with a Coomassie-blue-based reagent for visualization of the bound GST fusion protein (a representative example is shown in the lowest panel) and by X-ray film exposure for visualization of the bound radiolabeled protein (upper three panels). The migration of molecular weight markers is indicated on the right. Lanes 3 and 4 are technical replicates. (**D**) Quantification of the binding of human p110α, p85α and ERK2 proteins to GST or GST-WW. The results shown are the average of 5-21 independent repetitions of the of the type of binding assay shown in B and C (n = 12 for myc-tagged p110α, n = 21 for native p110α, n = 8 for p85α, n = 5 for ERK2). Error bars show the 95% confidence interval; individual data points are shown as dots. Technical replicates in a given experiment (e.g. lanes 3 and 4 in C) were averaged together to create a single data point. ****, p < 0.0001; ns, not significant (p > 0.05).

Initially, we used two different variants of full-length human p110α protein. The first version contained an N-terminal myc-epitope tag and a C-terminal CAAX box [38], and the second version was the unaltered 1086-residue wild-type protein. Quantification of 12-21 independent experiments for each variant is shown in Figure 1D, and demonstrates that the presence or absence of the myc and CAAX tags did not affect the binding of GSTWW to p110α. Furthermore, for both variants, their binding to GST-WW was statistically-significantly different from their binding to GST alone, with a very high level of confidence (p < .0001). The unaltered wild-type variant was used in the experiments shown in the remainder of the manuscript. Figure 1D also shows that for p85 and ERK2, both of which were assessed repeatedly over multiple different experiments, the trace binding to GSTWW was indistinguishable from the level of binding to GST alone (p > .99 for p85α, p > .40 for ERK2).

In addition to not binding to ERK2 (Figure 1), GST-WW also did not exhibit detectable binding to human ERK1, MEK1 or MEK2 (Figure 2 A,B). These negative results with RAS/MAPK pathway components confirmed our previously published data that used a different experimental approach to show that ERK1, ERK2, MEK1 and MEK2 all bind to the IQ domain of IQGAP1, and not to the WW domain [32]. In addition, these results demonstrated that our GST-WW construct was not non-specifically sticky.

**Figure 2.**
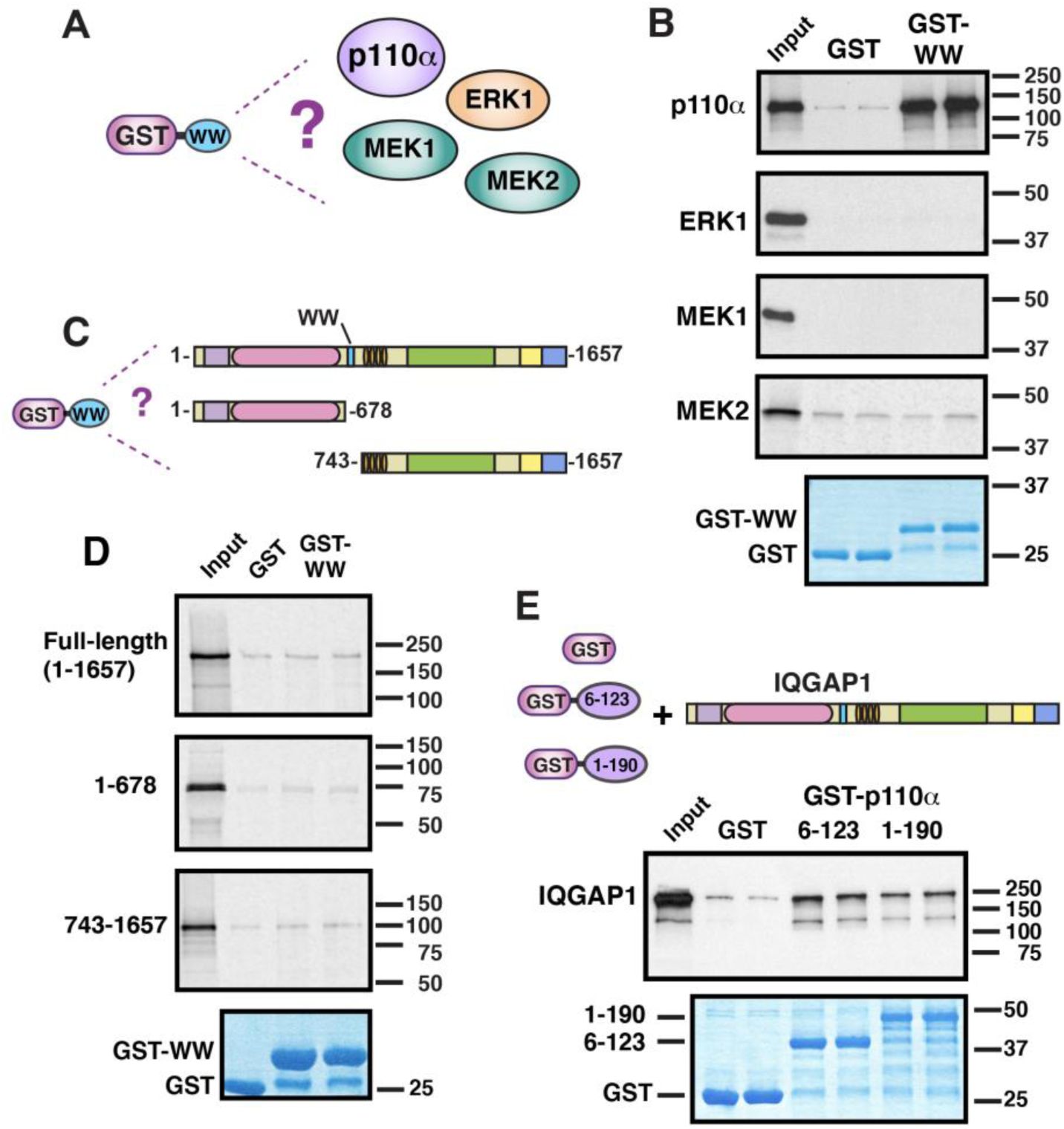
Additional binding and non-binding interactions involving IQGAP1 and its WW domain. (**A**) The WW domain of human IQGAP1, fused to GST, was tested for binding to full-length human p110α, ERK1, MEK1 and MEK2. (**B**) The WW domain binds to p110α, but not to ERK1, MEK1, or MEK2. Autoradiogram of a representative experiment of binding assays described in **A**. Each binding assay shown was repeated three separate times (i.e. three independent experiments), with duplicate points (i.e. technical replicates) in each experiment. Other details as in Figure 1. (**C**) The WW domain of human IQGAP1, fused to GST, was tested for binding to full-length human IQGAP1, as well as to the 1-678 and 743-1657 fragments of IQGAP1. (**D**) The WW does not bind to IQGAP1 *in trans*. Autoradiogram of a representative experiment of binding assays shown in **C**. Other details as above. (**E**) The adapter-binding domain of p110α binds to full-length IQGAP1. Residues 6-123 and 1-190 of human PIK3CA/p110α, fused to GST, were tested for binding to ^35^S-labelled, full-length human IQGAP1. Other details as above.

The WW domains of some proteins bind to other regions of the same protein; these in-trans or in-cis interactions mediate homodimerization or autoinhibition [39, 40]. Hence, we also asked whether GST-WW was able to bind to full-length IQGAP1 protein, or to IQGAP1 truncations lacking the WW domain. We found no evidence for such self-interaction (Figure 2 C,D). However, it should be noted that our binding assay would not be able to detect weak, millimolar-affinity interactions that might nevertheless be significant if driven by local concentration effects.

Preliminary domain-mapping experiments indicated that the N-terminal adapter-binding domain (ABD) of p110α was sufficient for binding to the WW domain of IQGAP1 (data not shown; these data will be published in a forthcoming study). In order to ask if full-length IQGAP1 protein could bind to p110α, we expressed two different fusions of the ABD to GST, and purified them from bacteria. GST-p110α(6-123) encodes the core of the ABD, whereas GST-p110α(1-190) includes N-terminal residues (residues 1-5) that are not visible in the crystal structure, as well as C-terminal residues (residues 124-190) that the link the ADB to the Ras-binding domain [41]. As shown in Figure 2E, both of these fusion proteins bound to full-length IQGAP1 protein that had been produced by *in vitro* translation. Hence, the WW domain of IQGAP1, purified from bacteria, binds to full length p110α (Figure 1, Figure 2A,B), and the adapter-binding domain of p110α, purified from bacteria, binds to full length IQGAP1 (Figure 2E).

To summarize, the experiments described in this section demonstrate that the WW domain of human IQGAP1 binds directly and specifically to the catalytic subunit of human PI 3-kinase.

### The cell-penetrating anti-tumor WW peptide specifically binds to p110α *in vitro*

As mentioned in the Introduction to this paper, a cell-penetrating IQGAP1 WW peptide has shown an ability to selectively kill various neoplastically transformed cells and primary clinical specimens in culture and to inhibit tumorigenesis and prolong survival in mouse models [12, 22, 27, 28]. As originally designed by Jameson *et al*. [12], this peptide contains, in order from N-terminus to C-terminus, eight D-arginine residues to promote cell penetration, a 10-residue myc-epitope tag to facilitate detection by immunostaining, and residues 680-711 of human IQGAP1. Some workers have raised the concern that the addition of the polyarginine or epitope sequences may have altered the binding specificity of this peptide [42]. To begin to explore this issue, we synthesized the identical peptide, and analyzed its *in vitro* binding specificity (Figure 3). As seen with GST-WW purified from bacteria (Figures 1 and 2), the synthetic anti-tumor WW peptide bound to p110α, but did not exhibit detectable binding to p85α, ERK1 or ERK2 (Figure 3A, B). Thus, the poly-D-arginine and myc-epitope sequences did not alter the binding specificity of the WW domain with the ligands tested. In particular, they did not endow the WW domain with an ability to bind to ERK1 or ERK2.

**Figure 3.**
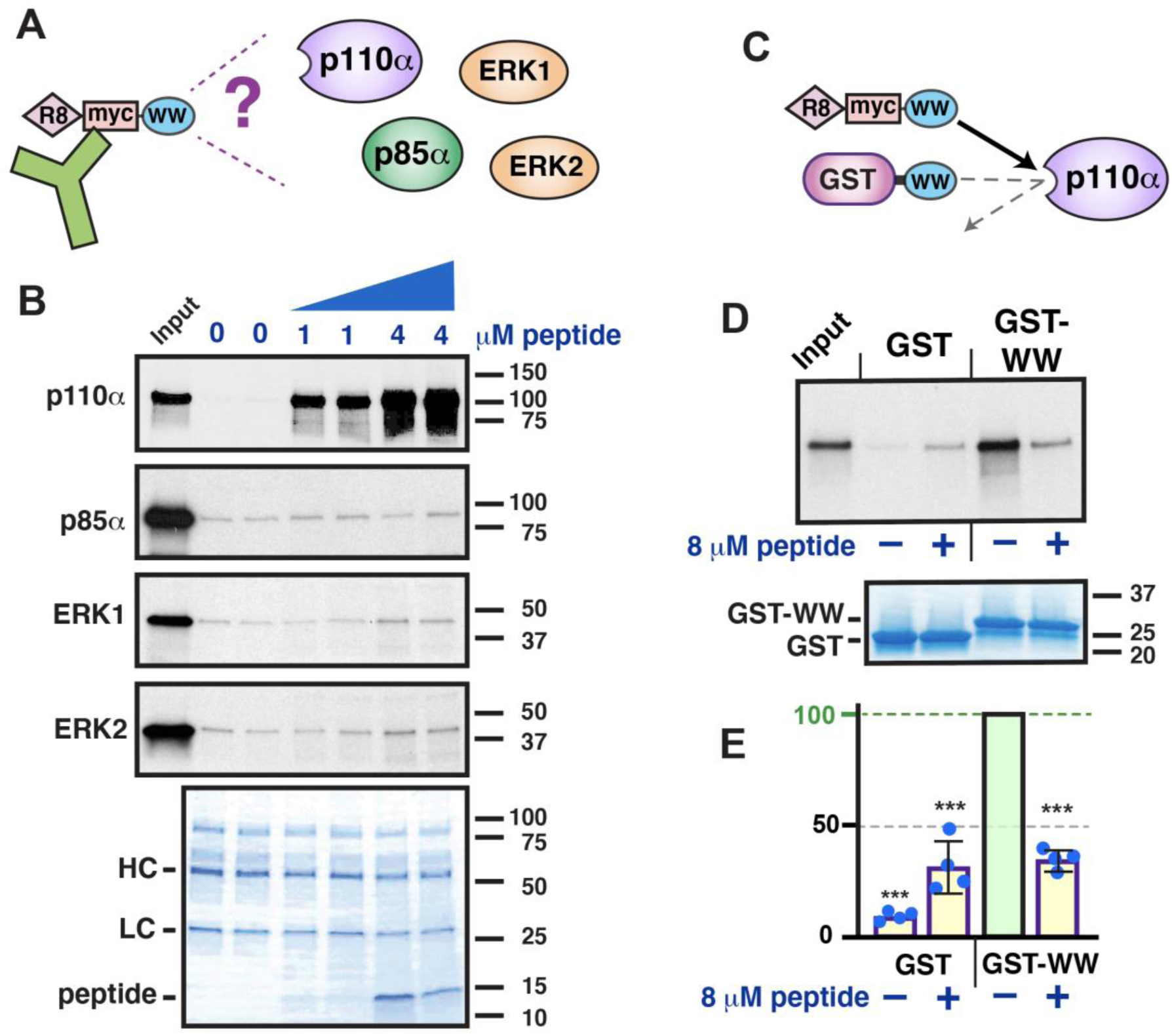
Binding specificity of the anti-tumor WW peptide. (**A**) The 50-residue anti-tumor WW peptide contains a poly-D-Arg motif (‘R8’), a myc epitope (‘myc’), and the WW domain of human IQGAP1 (‘WW’). The peptide was tested for its ability to coimmunoprecipitate p110α, p85α, ERK1 or ERK2. (**B**) The WW peptide binds to p110α but not to p85α, ERK1 or ERK2. As shown in **A**, ^35^S-radiolabeled full-length human p110α, p85α, ERK1 and ERK2 proteins were incubated with 0, 1 or 4 μM of WW peptide. The mixture was then incubated with an anti-myc-epitope antibody bound to protein A/G agarose beads, and the resulting bead-bound protein complexes were isolated by sedimentation and resolved by 4-20% SDS-PAGE. The gel was analyzed by staining with a Coomassie-blue-based reagent for visualization of the antibody heavy chain (HC) and light chain (LC); the peptide was also visible. The top four panels are X-ray film exposure of the bound radiolabeled protein. The p85α, ERK1 and ERK2 gels were overexposed to highlight the absence of any binding above the trace amount seen with the GST control. The amount in the ‘Input’ lanes is 5% of that added to the binding reactions. The migration of molecular weight markers is indicated on the right. (**C**) The WW peptide was tested for its ability to competitively inhibit the binding of GST-WW to p110α. (**D**) The WW peptide competes with GST-WW for p110α binding. As shown in **C**, ^35^S-radiolabeled fulllength human p110α protein was incubated with 25 μg (∼ 4 μM) GST-WW bound to glutathionesepharose beads, in the presence (+) or absence (–) of 8 μM WW peptide. Bead-bound complexes were then isolated and analyzed as described in Figure 1. (**E**) Quantification of four independent repetitions of the competition assay shown in **D**. Experiments were normalized by setting the binding to GST-WW in the absence of competing peptide to 100%. Error bars show the 95% confidence interval; individual data points are shown as dots. Significance estimates are based on the 95% confidence interval; see Experimental Procedures for details. ***, difference from normalized control is very highly significant.

We also tested the ability of the WW peptide to act as a competitive inhibitor of p110α binding. As shown in Figure 3 C and 4D, the WW peptide was indeed able to effectively compete with GST-WW for binding to p110α.

### The WW domain of IQGAP1 binds to the p85α/p110α complex

In cells, the majority of p110α molecules are bound to p85α in the form of p110α/p85α heterodimers, although p85α also has independent functions [43]. To ask if the WW domain of IQGAP1 could bind to p110α/p85α heterodimers, we co-translated p110α and p85α in the same *in vitro*-translation reaction, and then assessed the binding of this cotranslated mix to GST-WW; p110α translated alone and p85α translated alone were included for comparison (Figure 4A). As shown in Figure 1 and in Figure 4B, p85α alone did not display detectable binding to GST-WW, whereas p110α bound efficiently. Nevertheless, p85α co-translated with p110α bound to GST-WW (Figure 4B, see also Supplementary Figure 2). Indeed, the results, when quantified and corrected for labeling efficiency, indicated that there was an approximately 1:1 molar ratio of p85α to p110α in the bound fraction (see Figure 4B legend and Supplementary Figure 2 for quantification). This experiment demonstrates that the WW domain of IQGAP1 is able to bind to p110α/p85α heterodimers as well as to p110α monomers.

**Figure 4.**
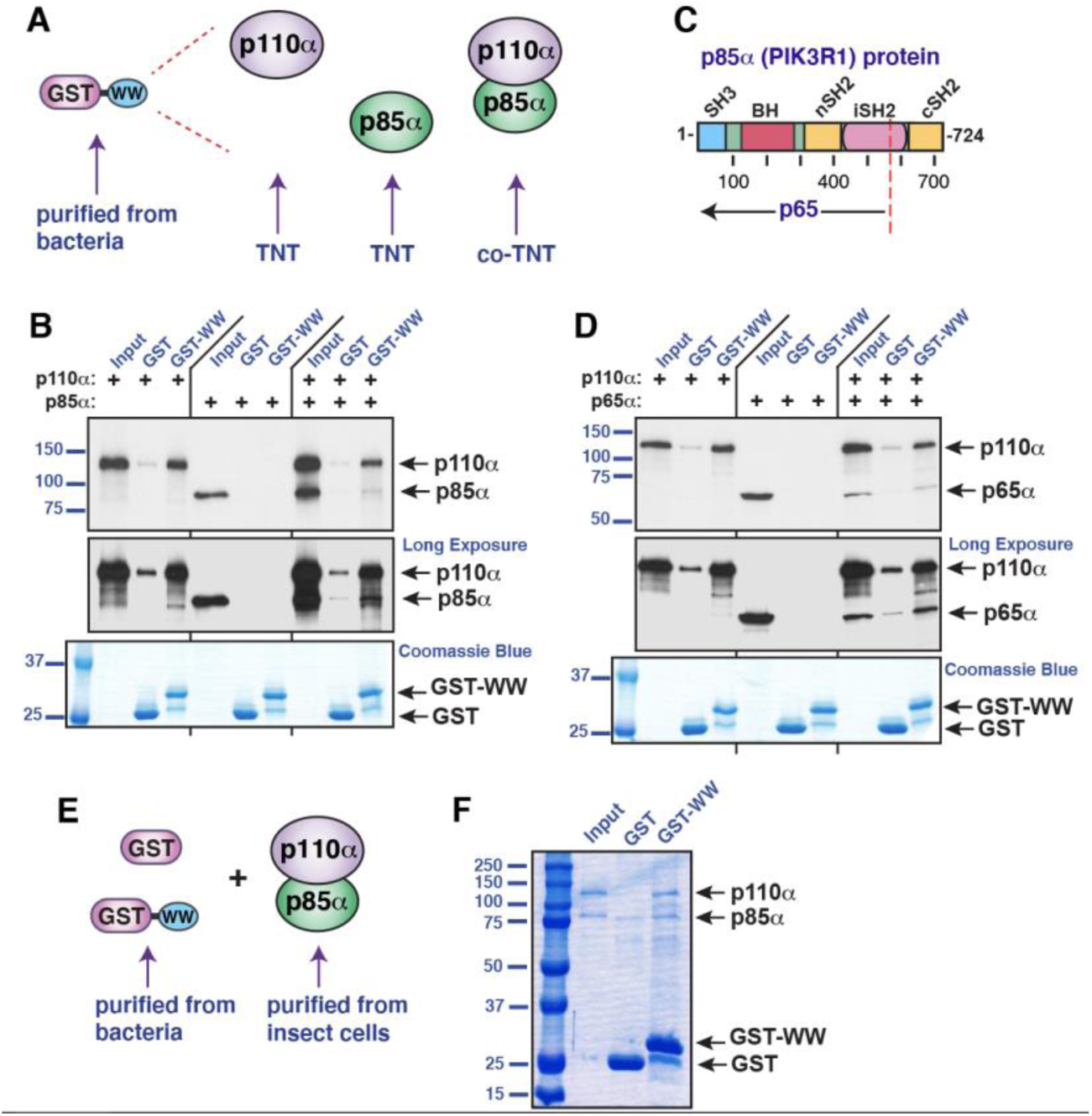
The WW domain of IQGAP binds to the PI 3-kinase holoenzyme. (**A**) Human p110α and human p85α were produced by coupled *in vitro*-transcription and translation (TNT) either separately or in the same reaction (“co-TNT”), and then tested for binding to GST or GSTWW. (**B**) Autoradiogram of a representative experiment of binding assays described in **A**. Top panel, standard exposure; middle panel, longer exposure; bottom panel, Coomassie-blue-based staining to visualize GST and GST-WW. Each binding assay shown was repeated three separate times (i.e. three independent experiments), with duplicate points (i.e. technical replicates) in each experiment. Other details as in Figure 1. After adjusting for labeling efficiency, the proportion of sedimented p85α/p110α was 1.47±0.87 (mean±95% confidence interval, n = 7); the 95% confidence interval thus includes the expected ratio of 1.0. (**C**) Schematic depicting full-length human p85α protein with domains denoted as follows: SH3, Srchomology 3 domain; BH, BcR homology domain; nSH2, N-terminal SH2 domain; iSH2, coiled coil region between the two SH2 domains; cSH2, C-terminal SH2 domain. The position of the truncation mutation that creates the activated p65α is shown by a dashed red line. (**D**) Human p110α and human p65α were produced by coupled *in vitro*-transcription and translation (TNT) either separately or in the same reaction, and then tested for binding to GST or GST-WW. Other details as in **B**. After adjusting for labeling efficiency, the proportion of sedimented p65α/p110α was 1.28±0.24 (mean±95% confidence interval, n = 8); the 95% confidence interval thus includes the expected ratio of 1.0. (**E**) Recombinant PI3K holoenzyme purified from insect cells was tested for binding to GST or GST-WW purified from bacteria. (**F**) 10 μg of purified holoenzyme was incubated with 25 ug of GST or GST-WW bound to glutathione Sepharose beads. The resulting bead-bound protein complexes were isolated by sedimentation and resolved by 10% SDS-PAGE. 5% (0.5 μg) of the holoenzyme in the binding assay points was loaded in the ‘Input’ lane. The gel was analyzed by Coomassie blue staining.

In the p110α/p85α heterodimer, the catalytic activity of p110α is repressed by p85α, but this repression is relieved when the two SH2 domains in p85α bind to phosphotyrosine residues on receptor tyrosine kinases or adaptor proteins (see Figure 4C for the domain structure of p85α). This repression can also be relieved by mutations in p110α or p85α [44, 45]. For p85α (gene = *PIK3R1*), such mutations cluster in the iSH2 domain, and are often nonsense or frameshift mutations that result in the expression of a shortened version designated p65α [46–48]. In order to determine if the WW domain of IQGAP1 could bind to the mutationally activated p110α/p65α heterodimer, we co-translated p110α with a truncated derivative of *PIK3R1* encoding p65α (residues 1-571, Figure 4C); p110α translated alone and p65α translated alone were also included for comparison. As shown in Figure 4D, p65α alone did not display detectable binding to GST-WW, whereas p110α once again bound efficiently. However, similar to the results obtained with p85α, p65α co-translated with p110α bound to GST-WW, with an approximately 1:1 molar ratio of p65α to p110α in the bound fraction (Figure 4D, see also Supplementary Figure 2).

To further demonstrate that p110α/p85α heterodimer can bind to the WW domain of IQGAP1, we obtained recombinant PI3K holoenzyme purified from insect cells, and assessed binding to GST-WW purified from bacteria (Figure 4E). As shown in Figure 4F, both the p110α and p85α subunits of PI3K co-sedimented with GST-WW, confirming that the purified holoenzyme does indeed bind to GST-WW. In addition, this experiment rules out the possibility that other macromolecules (such as other proteins, DNA, or RNAs) present in the rabbit reticulocyte lysate used to produce radiolabelled proteins for this study are necessary for the WW-p110α interaction.

To summarize, the results in this section show that the WW domain of IQGAP1 is able to bind directly to both the unactivated p110α/p85α heterodimer and the mutationally activated p110α/p65α heterodimer.

### The WW domain of IQGAP1 associates with PI3K in human HEK293 cells

To determine if the WW-binding domain of IQGAP1 was sufficient to associate with PI3K in cells, a plasmid expressing GST-WW was transiently transfected into the human HEK293 cell line, cell extracts were prepared, and GST-WW protein and associated proteins were isolated by sedimentation. As shown in Figure 5, both p110α and p85α were co-sedimented with GST-WW, but not with the GST alone control (Figure 5, lower panels, lanes 1 and 2).

**Figure 5.**
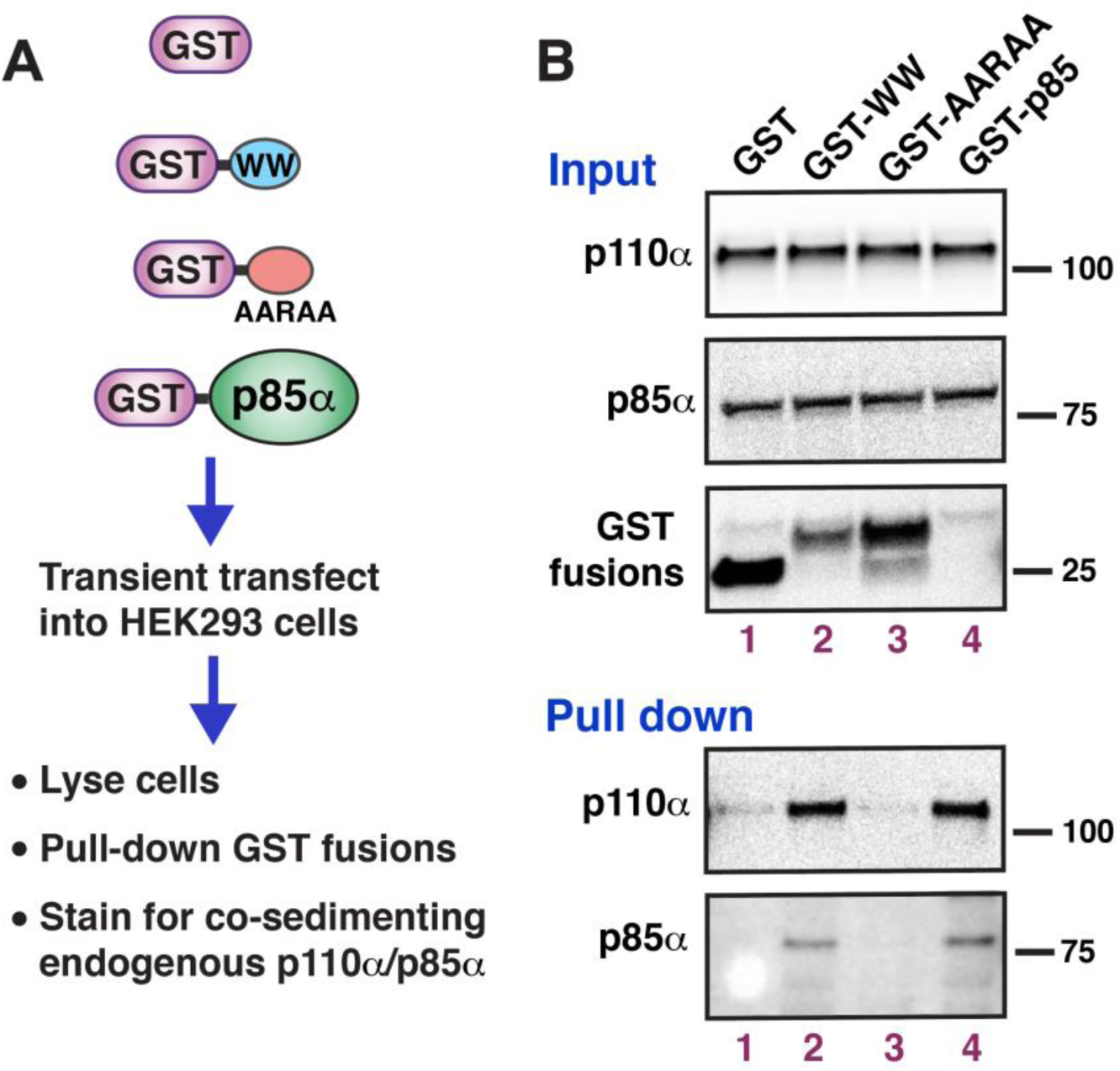
The WW domain of IQGAP1 associates with the PI3K holoenzyme in mammalian cells. (**A**) The constructs shown were separately transfected into HEK293 cells. The cells were then lysed, GST fusion proteins were isolated by sedimentation, and the sedimented samples were analyzed for associated p110α/p85α by immunoblotting. (**B**) Upper 3 panels: 20 μg of total cell lysates expressing the constructs shown in **A** were run on a 4-12% SDS PAGE gradient gel and immunoblotted with anti-p110α or anti p85α antibodies to determine the endogenous level of these proteins, and with anti-GST to assess the expression of the transfected GST fusion. Lower two panels: the sedimented fractions were analyzed for co-sedimented p110α and p85α by immunoblotting.

In the same experiment, we expressed two additional controls. The first of these was GST-AARAA, a mutant variant of IQGAP1’s WW domain that contains multiple substitutions in key conserved residues; as will be shown in Figure 7, this variant does not bind to p110α *in vitro*. This mutant also failed to asociate with p110α in cells (Figure 5, lane 3).

**Figure 6.**
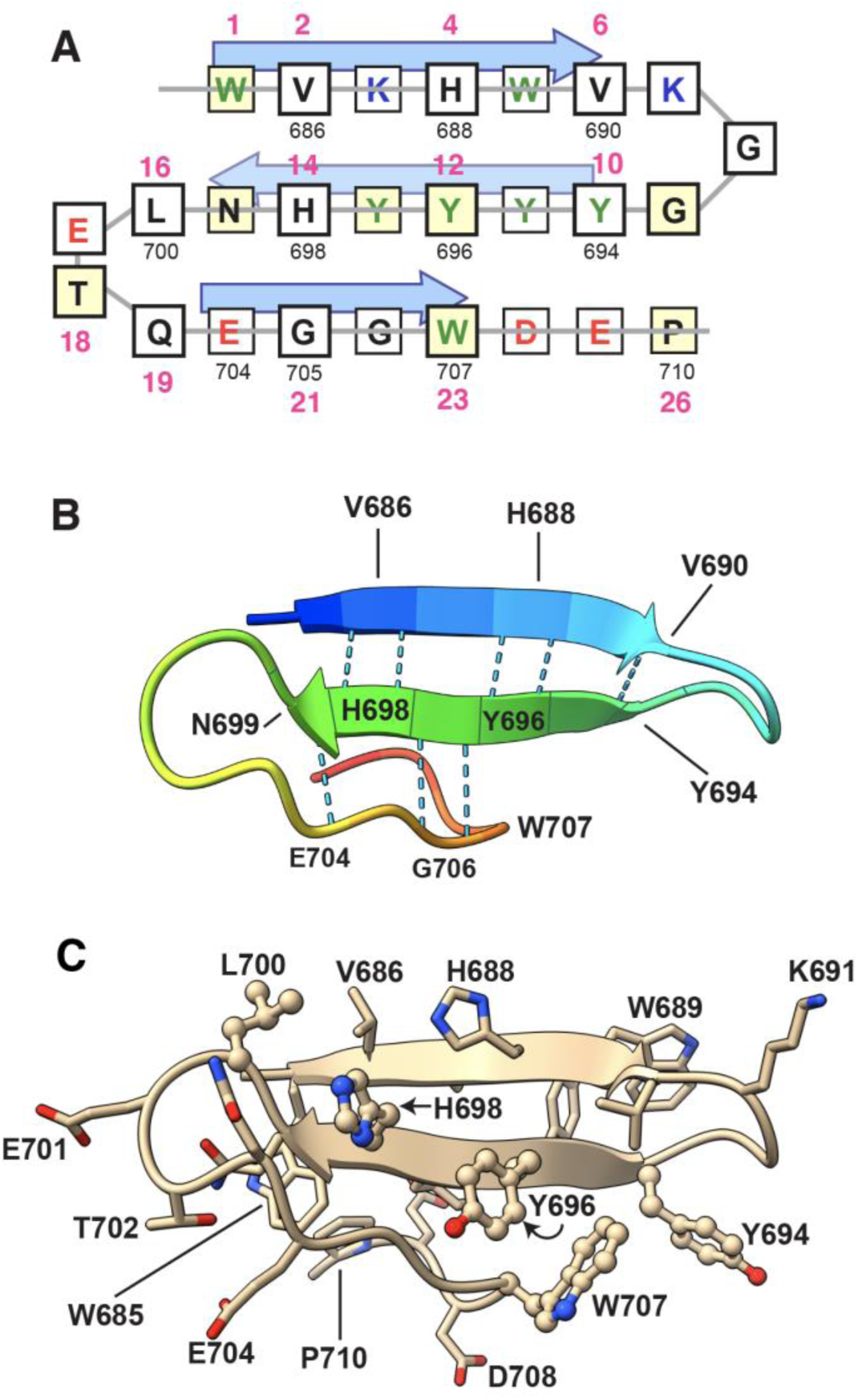
Model of the WW domain of human IQGAP1. (**A**) Schematic model. Gray lines show the chain tracing, amino acid residues are shown boxed in the single-letter code. Aromatic residues (W, Y, F) are green; positively-charged residues (R, K) are blue; negatively-charged residues (D, E) are red. Residues predicted to point up are above the gray line and have slightly larger boxes with thicker outlines. Yellow boxes indicate IQGAP1 residues that are conserved in most other WW domains. Arrows show the predicted β-strands. The smaller black numbers are the residue numbers in the 1657-residue IQGAP1 protein. The larger pink numbers are a relative numbering system wherein the first domain-defining tryptophan of the WW domain is designated position 1. (**B**) Ribbon diagram derived from the Alphafold-predicted structure, colored using the rainbow coloring option of the ChimeraX software package. Main-chain hydrogen bonds between adjacent β-strand residues are shown as dotted lines; other hydrogen bonds involving side chains are not shown. (**C**) Ribbon diagram derived from Alphafold-predicted structure with side chains shown. Side chain oxygen atoms are red, side chain nitrogen atoms are blue. The five residues subjected to single-site mutational analysis in the experiments shown in Figure 7 are shown in ball and stick format.

**Figure 7.**
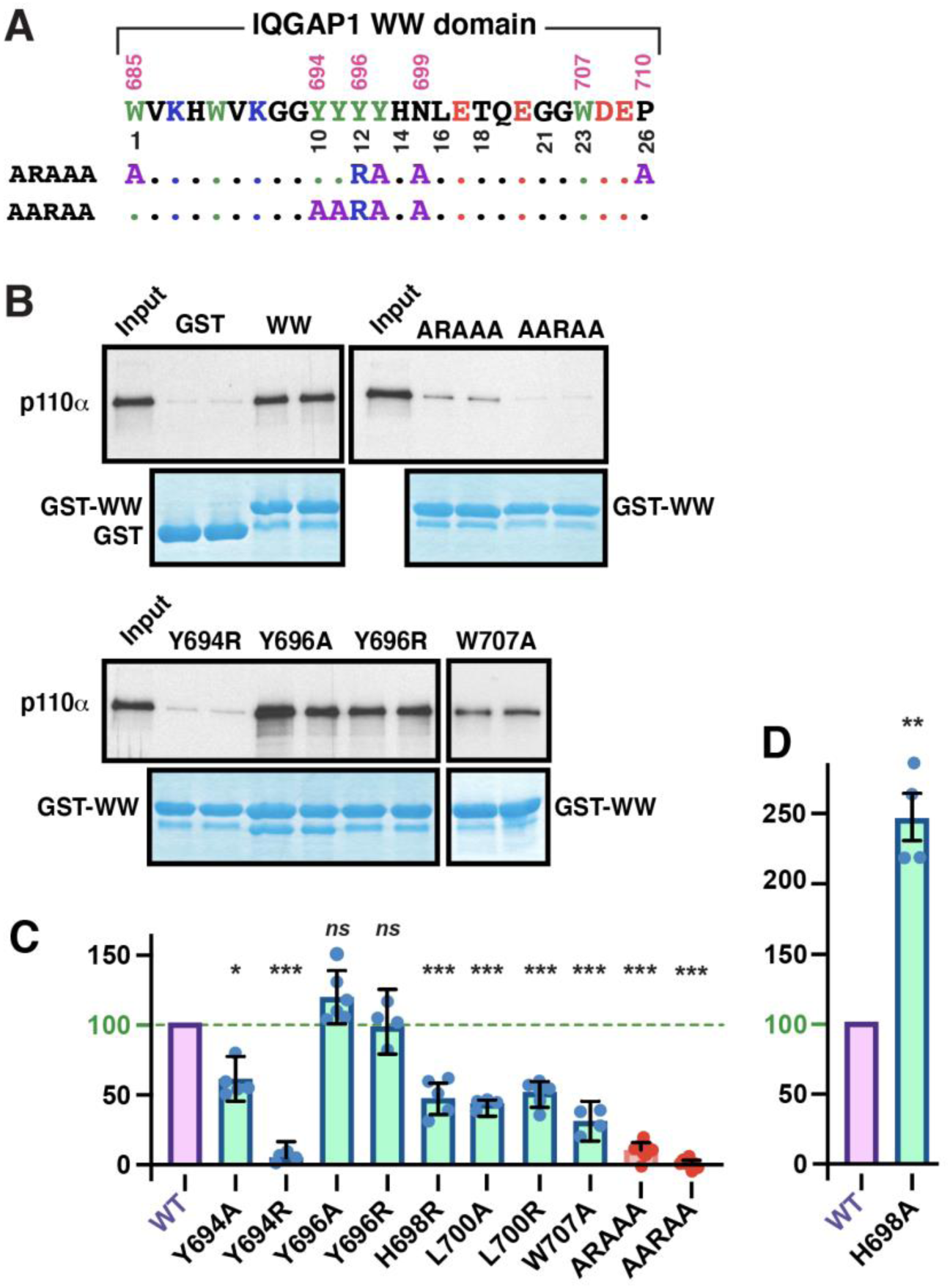
Structure/function analysis of the IQGAP1 WW domain by site-directed mutagenesis. (**A**) The top line shows the sequence of the WW domain of human IQGAP1. Residue numbers are shown above; the numbers below are a relative numbering system described in the main text. Below that, the sequence changes in two quintuplely-mutated IQGAP1 WW domain variants used in this study are shown. (**B**) The binding of GST, GST-WW and GST-WW variants to radiolabeled p110α was assessed. Autoradiograms of representative experiments of select variants are shown. (**C, D**) Quantification binding of GST, GST-WW and GST-WW variants to radiolabeled p110α, normalized to the percent binding of wild-type GST-WW. The results shown are the average of 4-8 independent repetitions, with duplicate points (i.e. technical replicates) in each repetition. Error bars show the 95% confidence interval (n = 4-8). The scatter of the individual data points is also shown. Significance estimates are based on the 95% confidence interval; see Experimental Procedures for details. *ns*, not significantly differ from 100%; *, significant; **, highly significant; ***, very highly significant.

The second control we expressed was GST-p85α, which we anticipated would serve as a positive control for p110α binding. This protein was very weakly expressed (data not shown), yet its presence was evident by its ability to pull-down endogenous p110α from the extract (Figure 5 lane 4, lower panels). Endogenous p85α also co-sedimented with GST-p85α; this is presumably attributable to the ability of p85α to homodimerize [49, 50]. To summarize, the WW domain of IQGAP1 specifically associated with p110α/p85α in HEK293 cells.

### A model of IQGAP1’s WW domain

WW domains are found in plants, fungi, protists, and animals. An alignment of the WW domain of IQGAP1 with 93 other well-recognized human WW domains, a tree, sequence logos, and other analyses are presented in Supplementary Figures 3–6. These analyses revealed that the WW domains closest in sequence space to IQGAP1 include the single WW domains from IQGAP3, IQGAP2 and WAC, as well as both of the tandem WW domains from WBP4 (a.k.a. FBP21), PRPF40A (a.k.a. FBP11), PRPF40B, TCERG1 (a.k.a. FBP28) and ARHGAP12. There are no published or unpublished structures of the WW domain from any IQGAP ortholog in the Protein Data Bank. Nevertheless, by comparing the sequence of IQGAP1’s WW domain to published structures of WW domains from WBP4, TCERG1 and PRPF40A [51–55], we constructed a schematic model of IQGAP1’s WW domain (Figure 6A). Human IQGAP has 21 residues between the two domain-defining tryptophans, a feature shared with over 80% of human WW domains. Thus, henceforth, we will use the numbering scheme shown in Figures 6A and 7A, in which these tryptophans are at positions 1 and 23, to discuss general sequence features of WW domains. A tryptophan at position 1 and a tryptophan or phenylalanine at position 23 are found in over 95% of human WW domains, including IQGAP1’s. Other sequence features that IQGAP1 shares with the majority of human WW domains include a glycine at position 9, aromatic residues at positions 12 and 13, an asparagine or aspartate at position 15, a threonine position 18, and a proline at position 26.

The schematic representation (Figure 6A) is consistent with the structure predicted by Alphafold [56, 57], which became available subsequently (Figures 6B and 6C). In this structure, the WW domain of IQGAP1 is predicted to fold into a canonical WW domain structure: an antiparallel beta sheet with three short strands connected by two loops, with main-chain hydrogen bonds providing stability (Figure 6B). As in other WW domains [52, 53, 55, 58], the side chain of the first domain-defining tryptophan (residue 1 in Figure 6A, W685 in IQGAP1) faces the underside of the sheet, where, together with the tyrosine at position 13 (Y697 in IQGAP1), the asparagine at position 15 (N699 in IQGAP1) and the proline at position 26 (P710 in IQGAP1), it constitutes a hydrophobic core that provides additional stability to the domain (Figure 6C). Alternating side chains in the beta strands (residues 2, 4, 10, 12, 14, 16, 18, 21 and 23) face up relative to the plane of the sheet to form the top side of domain; these residues, as well as residues in one or both of the connecting loops, are frequently observed to participate in ligand binding in other WW domains [53–55, 59–62]. Collectively, these features suggest that the WW domain of IQGAP1 adopts a canonical WW domain structure [23–25]. There are, however, several differences between IQGAP1 and the consensus WW domain sequence that may be important (see further below and Discussion).

### Conserved hydrophobic core residues in the WW domain are necessary for p110α binding

Previously, we constructed a mutant version of IQGAP1 that contained five different substitution mutations in highly conserved residues in its WW domain [32]. These 5 changes include (i) alanine substitutions to four residues that constitute a hydrophobic core on the underside of the beta sheet in other WW domains [52, 53, 55, 58], and in the predicted structure shown in Figure 6; (ii) A tyrosine-to-arginine change in a residue (position 12/Y696 in Figure 7A) whose side chain points up into the ligand binding site and helps form the proline-binding groove in other WW domains [23, 53, 55]. In our previous work, we demonstrated that these 5 mutations did not influence the binding affinity of IQGAP1 for ERK2, consistent with our finding that the IQ domain of IQGAP1 was necessary and sufficient for binding to ERK2 [32]. Here, we reintroduced those 5 mutations (W685A, Y696R, Y697A, N699A, P710A) into GST-WW and renamed this variant ‘ARAAA’ (see Figure 7A). We also constructed a second quintuple mutation variant of GST-WW designated ‘AARAA’. This variant contains three of the above changes, plus two more substitutions to residues in the second beta strand. As shown in Figure 7B (top panels), both of these variants were completely defective for binding to p110α. The results of eight independent experiments with these two mutants is quantified in Fig. 7C, and indicated that the lack of binding exhibited by these two variants is very highly significant. Hence, combining multiple substitutions in conserved residues in the WW-domain of IQGAP1 abolished its ability to bind to p110α.

### Key conserved residues in the WW domain are necessary for p110α binding

To begin to explore residues in IQGAP1’s WW domain that may directly contact p110α, we used site-directed mutagenesis to examine the role of five individual residues thought to participate in ligand binding in other WW domains [51–55, 59–67]; these residues are represented in ball-and-stick format in Figure 6C. Moving from N-terminal to the C-terminal, the first residue we examined was position 10/Y694. This position, which sits at the beginning of beta strand 2, is an aromatic residue in only 4 other human WW domains; instead, most human WW domains have an arginine or lysine at this position. As shown in Figure 7B and 7C, substitution of Y694 with alanine reduced the binding of GST-WW binding to p110α by approximately 40%, and substitution with arginine abolished binding.

The next residue we examined is position 12/Y696. This residue is a tyrosine or phenylalanine in 96% of human WW domains, and its mutation to alanine [66] or arginine [64] greatly reduced ligand binding in other WW domains tested. In published structures, it is typically observed to help form the binding groove for one or more prolines in the ligand [23, 53, 55]. Interestingly, mutation of this residue to alanine had no detectable effect on GST-WW binding to p110α, nor did the more drastic substitution of arginine for the native tyrosine (Figure 7B, 7C).

In most human WW domains, the position 14 residue is usually either a branched-chain hydrophobic residue (valine, leucine or isoleucine; 63% of human WW domains) or an aromatic residue (30%). In human IQGAP1, position 14 is a histidine. Position 14 has been shown to be involved in ligand binding in multiple WW domains [53, 55, 59–61], although in none of these examples was the position 14 residue a histidine. As shown in Figure 7C, in the IQGAP1 WW domain, a histidine to arginine substitution at position 14/H698 reduced binding by roughly 50%. Interestingly, substitution of this residue with alanine reproducibly increased binding to p110α, by an average of 250% (Figure 7D).

Position 16 has also been shown to be involved in ligand binding in multiple studies [59–62], although in all these studies the position 16 residue was a histidine, as it is in the majority (57%) of human WW domains. In human IQGAP1, position 16/L700 is a leucine. As shown in Figure 7C, substitution of L700 with either alanine or arginine both reduced binding to p110α by about 50%.

Finally, position 23/W707, like position 12/Y696, is typically observed to help comprise the proline-binding groove [23, 53, 55, 59]. A tryptophan to alanine substitution at this position in GST-WW reduced binding to p110α by roughly 75% (Figure 6C).

To summarize, mutational changes to four of the five individual residues we examined resulted in decreased binding to p110α, consistent with expectations based on studies of other WW domains. On the other hand, there were some possibly important differences and nuances; see Discussion for further analysis.

### The WW-p110α interaction has a modest affinity

Over the course of these studies, we performed over 20 independent, quantitative binding assay experiments between the IQGAP1 WW domain and p110α (Figure 1D). From these data we were able to obtain an estimate of the 130 μM for the dissociation constant (K_d_) of this interaction, with a 95% confidence interval of 105-153 μM (see Supplementary Table 1). Typically, WW domains that have affinities in this range are aided in binding to their ligands by additional interactions between the binding partners that occur outside of the WW domain; see Discussion for further analysis.

## Discussion

IQGAP is a multifunctional scaffold protein that has been implicated in many physiological processes. It has an evolutionarily ancient role in regulating cytoskeletal rearrangements in fungi and animals [68, 69]. In mammals, it also functions as a scaffold protein for the RTK/RAS/MAPK pathway and the PI3K pathway – two of the pathways most frequently mutated in cancer [70–72] – and additionally regulates several other cancer-relevant signaling pathways [17–21]. Much work still needs to be done to understand how IQGAP1 and other scaffold proteins coordinate signaling in localized subcellular regions depending on cell type and cell context, how this may go awry in disease, and how it might be exploited for new targeted therapies. Despite the challenges, continued progress in this area may open new opportunities for personalized medicine. A promising direction in this regard is blocking kinase-scaffold interactions as a therapeutic strategy for treating cancer and other diseases.

Here we searched for a binding partner for the WW domain of IQGAP1, motivated by recent findings that a cell-penetrating version of this domain has potent anti-tumor properties with minimal toxicity to non-transformed cells and tissues (see Introduction for further details). We found that this domain binds directly to p110α, the catalytic subunit of PI 3-kinase, which is one of the most frequently mutated proteins in human cancer. In contrast, this WW domain did not bind to ERK1, ERK2, MEK1, MEK2, or other parts of IQGAP1 (Figure 1, Figure 2). Moreover, we found the same set of binding specificities for the cell-penetrating anti-tumor WW peptide that contains poly-D-Arginine and myc-epitope tags in addition to the WW domain residues (Figure 3). Further, we found that the WW domain of IQGAP1 did not bind directly to p85α, the regulatory subunit of PI 3-kinase, but could bind to the p110α/p85α holoenzyme, both *in vitro* (Figure 4) and in human tissue culture cells (Figure 5). IQGAP1’s WW domain also bound to the p110α/p65α heterodimer, which contains a truncated, mutationally-activated variant of p85α (Figure 4), demonstrating that the WW domain of IQGAP was able to bind to both unactivated and mutationally-activated PI3K. Finally, we built a model of IQGAP1’s WW domain (Figure 6), and demonstrated the role of key residues in this domain in p110α binding (Figure 7). Our results provide new insights into IQGAP1-mediated scaffolding, and suggest that cell-penetrating WW peptides may exert their anti-tumor effects by disrupting the interaction of IQGAP1 with the catalytic subunit of PI 3-kinase.

### A model of IQGAP1’s WW domain and the role of key residues in ligand binding

No x-ray or NMR structures of IQGAP1’s WW domain have been published. Nevertheless, by comparison to published structures of WW domains that are close in sequence to IQGAP1, we built a model of IQGAP1’s WW domain (Figure 6). Using this model as a reference, we then tested the role of selected residues in this WW domain on p110α binding. First, we made multiple substitutions to residues that constitute the hydrophobic core on the underside of the domain, and have been shown in other WW domains to contribute to folding and stability. Consistent with expectations, both multisubstituted variants we tested were severely defective in binding to p110α (Figure 7). We also expressed one of these multi-substituted variants in human cells, and showed that it was defective in interacting with endogenous PI3K (Figure 5).

Next we tested the role of individual residues in IQGAP1’s WW domain that are predicted to be situated on the ligand binding surface (Figure 7). We found that mutations to residues in positions 10, 14, 16 and 23 (see Figure 7A for numbering) all affected the binding of GST-WW to p110α. These results are consistent with the hypothesis that these residues either directly contact p110α, or influence the positioning of residues that do.

The results of mutating position 16 (L700 in IQGAP1) and position 23/W707 were the most straightforward. All tested mutations of these residues reduced binding to p110α by 50% or more. Moreover, there is considerable precedent for a direct role in ligand binding for residues in these positions in other WW domains [23, 53, 55, 59–62].

The results with position 14/H698 were more complicated, in that substitution with alanine actually increased binding to p110α by about 250%, whereas substitution with arginine reduced binding by over 50%. Interestingly, position 14 is known to be a key residue in determining the specificity of binding in other WW domains [60]. Moreover, substitutions to the position 14 residue in the WW domain of PIN1 have also been shown to increase ligand binding affinity [73]. Modifications of H698 may thus be worth pursuing as a means to develop IQGAP1 WW peptides and peptidomimetics that have higher affinity for p110α, and therefore might display greater anti-tumor potency as well.

In many published structures of WW-ligand complexes, the position 10 residue is not observed to make direct contact with the ligand. In both cases where the position 10 residue does contact the ligand, position 10 is an arginine [65, 67]; indeed, the position 10 residue is a basic, positively charged arginine or lysine in most human WW domains. In contrast, position 10 in IQGAP1 is a tyrosine. We found that substituting this residue with arginine (Y694R) virtually eliminated binding to p110α, and that substitution with alanine reduced binding by about 40% (Figure 7). The simplest interpretation of these data is that Y694 participates directly in binding to p110α.

In most published structures of WW-ligand complexes, the position 12 residue is aromatic, as it is in IQGAP1, and is observed to make direct contact with a proline in the ligand [23, 53, 55]. Furthermore, mutation of this residue to alanine [66] or arginine [64] was found to substantially compromise binding in other WW domains tested. For IQGAP1’s WW domain, however, we found that substituting this residue with either alanine (Y696A) or arginine (Y696R) had no discernable effect on binding to p110α (Figure 7). The simplest interpretation of these data is that the side chain of Y696 does not make critical contacts with p110α. Collectively, the results with position 10/Y694 and position 12/Y696 suggest that there may be some differences between how the WW domain of IQGAP1 binds p110α and the precedent established by the majority of WW domain studies. Another feature which suggests that there may be unique aspects to the manner in which the WW domain of IQGAP1 binds its ligand(s) is the presence of a glycine at position 21. IQGAP3 is the only other human WW domain containing a glycine at this position; instead, 86% of human WW domains have a serine or threonine here, and these side chains have been shown to participate in ligand binding in many instances [53, 55, 61, 62].

WW domains are proline recognition domains that typically bind to proline-rich short linear motifs in their binding partners [26]. Further work will be required to map the binding site(s) on p110α that bind to the WW domain of IQGAP1. In a systematic study of 42 WW domains that focused on identifying ligands, the WW domain of IQGAP1 was found to be unfolded under the conditions studied, and a ligand could not be identified [63]. The lack of ligand identification in this previous study cannot be solely attributed to the lack of folding, since some WW domains are observed to fold in the presence of ligand [63, 74, 75]. If indeed there is coupled folding and binding in the interaction of IQGAP1’s WW domain with p110α, this could help explain the modest affinity of the interaction that we observed. It is a well-established hypothesis that coupled folding and binding may promote low-affinity yet high-specificity molecular recognition [76].

### An updated model of IQGAP1-mediated scaffolding interactions

It is difficult to express a PI3K catalytic subunit like p110α in the absence of a regulatory subunit such a p85α, or vis versa, as the expressed protein either forms heterodimers with its endogenous partner or is rapidly degraded [41, 43, 44]. We were able to overcome this difficulty by using a rabbit reticulocyte lystate *in vitro* translation system to express p110α and p85α. Reticulocytes do not express endogenous p110α/p85α [77], and we found no evidence of p110α or p85α degradation in this system, even when the proteins were translated separately.

Using this approach, we showed that the WW domain of IQGAP1 binds directly to p110α, yet we found no evidence that this WW domain binds directly to p85α. Anderson and colleagues, however, have shown that both the WW and IQ domains of IQGAP1 are necessary for its binding to the PI3K holoenzyme in cells, and that a fragment of IQGAP1 containing both of these domains binds directly to p85α [22]. A plausible model accounting for all these observations is that both the WW domain and the IQ domain of IQGAP1 cooperate to bind to the p110α/p85α holoenzyme, with the WW directly contacting p110α, and the IQ domain contacting p85α. This cooperation may function to increase the affinity of IQGAP1-PI3K binding, as well as to optimally position PI3K relative to a localized pool of its substrate PIP2. Cooperative ligand binding involving a WW domain and a neighboring domain has been observed in other cases [23]. For example, the WW domain in dystrophin cannot bind its ligand without the help of the adjacent EF hand domain [61]. Furthermore, in proteins that contain more than one WW domain, two such domains can cooperate to bind their target with higher affinity and specificity [78, 79].

Figure 8 shows a model of select binding and scaffolding interactions mediated by the WW and IQ domains of IQGAP1, based on pioneering work by other groups [12, 22, 29, 80–83], and extended by the findings presented here and in our previous study [32]. First, multiple components of the RTK/RAS/MAPK cascade –including RTKs, BRAF, MEK1/2, and ERK1/2– interact with the IQ domain of IQGAP1 (Figure 8A). Second, in the PI3K pathway, IQGAP1 interacts with PI3K using both its WW and IQ domains, the former recognizing p110α and the latter primarily recognizing p85α (Figure 8B). It is also possible that there are further interactions between IQGAP1 and p110α in addition to those mediated by the WW domain. IQGAP1 also interacts with PIPKI, the enzyme immediately upstream of PI3K, via its IQ domain (Figure 8B) [22, 83, 84]. Third, we posit that cell-penetrating IQGAP1 WW peptides may block the productive engagement of PI3K with IQGAP1 by directly interfering with WW-p110 binding (Figure 8C).

**Figure 8.**
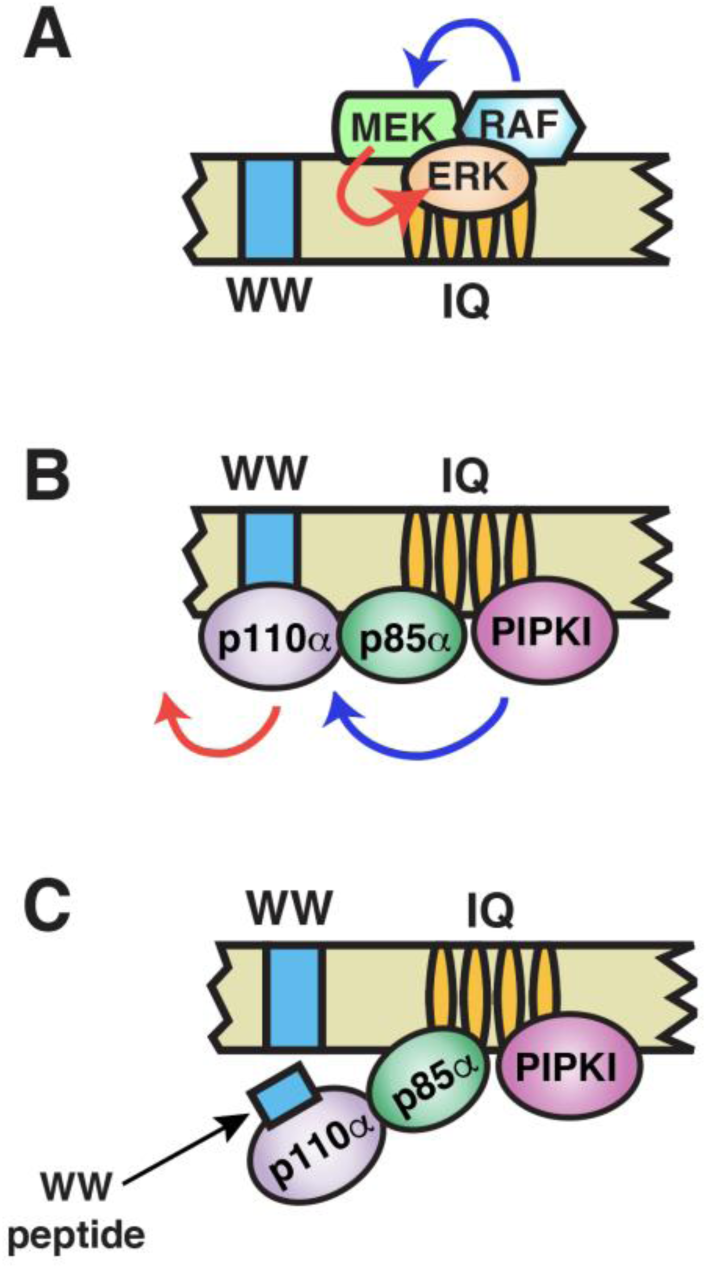
Model of key binding and scaffolding interactions mediated by the WW and IQ domains of IQGAP1 with elements of the MAPK and PI3K pathways. (**A**) MAPK cascade interactions. Both MEK and ERK bind to the IQ domain, in close proximity to RAF and some receptor tyrosine kinases, which also bind to this domain. These interactions may facilitate RAF phosphorylation of MEK, and MEK phosphorylation of ERK. (**B**) PI3K pathway interactions. PI3K binds to IQGAP1 using a bipartite mechanism: p110α binds to the WW domain, and p85α binds to the IQ domain. This may position PI3K in an optimal position and orientation to utilize a localized pool of its substrate PIP2. PIP2 is generated by PIPKI (a.k.a. PIP5K1A), which also binds to the IQ domain. (**C**) We propose that cell-penetrating anti-neoplastic WW peptides may block the productive engagement of p110α with the WW domain on IQGAP1. See text for further details.

### Summary and conclusions

Herein we showed that the p110α catalytic subunit of human phosphoinositide 3kinase (PI3K) is a direct binding partner of the WW domain of human IQGAP1, and made progress on the structure/function characterization of this interaction. Our findings propound a refined model of IQGAP1-mediated scaffolding, and suggest that blocking the productive engagement of PI3K with IQGAP1 is a key on-target effect of anti-neoplastic IQGAP1 WW-derived peptides.

## Experimental Procedures

### Genes

The mammalian genes used in this study were human *IQGAP1* (NCBI accession number NM_003870), bovine *PIK3CA* (NM_174574), human *PIK3CA* (NP_006209), human *PIK3R1* (NM_181523), human *ERK1* (MAPK3; NM_002746), human *ERK2* (MAPK1, NP_620407), human *MEK1* (MAPK2K1, NM_002755), and human *MEK2* (MAP2K2, NM_030662).

### IQGAP1-based constructs

GST-WW contains residues 679-719 of human IQGAP1 inserted into plasmid pGEXLB [85]. The region of interest was amplified in a polymerase chain reaction (PCR) using plasmid pCR-BluntII-TOPO-hIQGAP1 [32] as the template and primers hIQGAP1(679-x) and hIQGAP1(x-719) (See Supplementary Table 2 for all PCR primer sequences). The PCR product was cut with *BamH*I and *Sal*I and cloned into pGEXLB that had been digested with the same enzymes. For the experiment shown in Figure 3 and elsewhere, QuikChange (Agilent) was used for site-directed mutagenesis; all mutations were confirmed by sequencing.

For the experiments shown in Figure 2, the constructs for producing full-length IQGAP1 protein and IQGAP(1-678) by *in vitro* transcription/translation have been described [32]. To make IQGAP1(743-1657), the region of interest was amplified in a PCR using plasmid pCR-BluntII-TOPO-hIQGAP1 as the template and primers hIQGAP1(743– x) and hIQGAP1(x-1657). The PCR product was cut with *Xma*III and *Sal*I and cloned into pGEM3Z that had been digested with the same enzymes. The resulting construct, pGEM3Z-hIQGAP1(743-1657) contains the 743-1657 open reading frame downstream of a T7 promoter.

### PIK3CA and PIK3R1-based constructs

A plasmid containing the bovine *PIK3CA* gene with an N-terminal myc-epitope tag and a C-terminal CAAX box, downstream of a T7 promoter, was obtained as a gift from Julian Downward [38]. We converted this to a sequence that encodes human p110α protein by site directed mutagenesis, as the human and bovine amino acid sequences differ at only two residues. Specifically, we introduced R532K and C535S substitutions into the bovine coding sequence. The resulting plasmid is designated T7-hb-p110α, where ‘hb’ stands for “humanized bovine”.

Plasmid pGEM3Z-p110α contains the coding sequence for full-length untagged human p110a protein downstream of a T7 promoter. To construct this plasmid, the human p110α-encoding open reading frame was amplified by PCR using primers p110-1-x and p110-end-down and T7-hb-p110α as the template. This process removed both the myc tag and the CAAX box, and replaced them with wild-type sequence. The resulting PCR product was digested with *BamH*I and *Sal*I and inserted into the corresponding sites of pGEM3Z.

Plasmid pGEM3Z-p85α contains the coding sequence for full-length human p85α protein downstream of a T7 promoter. To construct this plasmid, the *PIK3R1* open reading frame was amplified by PCR using primers p85up and p85down. The template for this reaction was a human *PIK3R1* sequence-verified cDNA clone from the mammalian gene collection and sold by Dharmacon (Clone ID 30528412). The resulting PCR product was digested with *BamH*I and *Sal*I and inserted into the corresponding sites of pGEM3Z.

Plasmid pGEM3Z-p65α (encoding residues 1-571 of p85α protein) was constructed from pGEM3Z-p85α by introducing a stop codon at codon 572 of the *PIK3R1* coding sequence by site-directed mutagenesis.

To construct GST-p110α(1-190) GST-p110α(6-123) for use in Figure 2E, the regions of interest were amplified in a polymerase chain reaction (PCR) using plasmid pGEM3Z–p110α as the template and primer p110-1-x or p110-6-x as the upstream primer and p110–x-190 or p110-x-123 as the downstream primer (Supplementary Table 2). The PCR product was cut with *BamH*I and *Sal*I and cloned into pGEXLB that had been digested with the same enzymes.

### ERK and MEK constructs

The constructs for *in vitro* transcription/translation of full-length ERK1, ERK2, MEK1 and MEK2 used in Figures 1-3 have been described elsewhere [85]. The MEK1(1-60) and MEK2(1-64) constructs used in Supplementary Figure 1 have also been described elsewhere [86].

### Constructs for mammalian cell expression

The constructs for the cell-based experiments in Figure 5 were made in multiple steps. First, the open reading frames encoding GST, GST-WW and GST-WW-AARAA were amplified in a PCR reaction using primers GST-WW-up and GST-WW-down (Supplementary Table 2) and the corresponding pGEXLB-based plasmid as the template. This strategy placed a unique *EcoR*I site upstream of the start codon for GST and a unique *Xba*I site downstream of the stop codon of the open reading frame (ORF). The PCR products were digested with *EcoR*I and *Xba*I and inserted into the mammalian expression vector pcDNA-V5-HIS-B (Invitrogen Corp.) that had been cut with the same enzymes. The resulting plasmids (pcDNA-GST, pcDNA-GST-WW, and pcDNA-GST-WW-AARAA) express the GST-fusion constructs driven by the CMV promoter; the V5 and HIS6 tags are not included in the fusion proteins.

pcDNA-GST-p85α was constructed by excising the human p85α ORF from pGEM3Z–p85α as a *BamH*I-*Sal*I fragment, and inserting into pGEXLB that had been cut with the same enzymes, yeilding pGEXLB-p85α, which served as a cloning intermediate. In the next step, the GST-p85α ORF in this plasmid was amplified using primer GST-WW-up and GST-WW-down and subcloned into pcDNA-V5-HIS-B as described above.

### *In vitro* transcription and translation

Proteins labeled with [^35^S]-methionine were produced by coupled transcription and translation reactions (T7, Promega). Translation products were partially purified by ammonium sulfate precipitation [87], and resuspended in binding buffer (20mM Tris-HCl (pH 7.1), 125mM KOAc, 0.5mM EDTA, 1mM DTT, 0.1% (v/v) Tween20, 12.5% (v/v) glycerol) prior to use in binding assays.

### Protein purification and binding assays

GST fusion proteins were expressed in bacteria, purified by affinity chromatography using glutathione-Sepharose (Cytiva Life Sciences) and quantified as described elsewhere [85, 88].

Protein binding assays were performed as described previously [85, 88]. Quantification of binding was performed on a Typhoon TRIO+ Imager using phosphorimaging mode. Percent binding was determined by comparing the input with the amount that was co-sedimented. This was then corrected for minor differences in loading of the GST fusion protein, which was assessed by quantifying a TIFF image of the Coomassie-blue-stained gel using ImageJ software. Each binding assay presented in this paper was repeated at least four separate times (i.e. four independent experiments), with duplicate points (i.e. technical replicates) in each experiment. Technical replicates in a given experiment are averaged together to obtain a single data point. We define ‘independent experiments’ as experiments performed on different days, with fresh batches of GST-fusion proteins and *in-vitro* translated proteins.

Co-immunoprecipitation experiments (Figure 3) were performed as described previously [89]. PIK3CA/PIK3R1 (p110a/p85a) recombinant human protein for use in the experiment shown in Figure 4F was purchased from Thermo Fisher Scientific (catalog number PV4788).

### Cell culture, transfection and pull-downs from human cells

Human embryonic kidney (HEK) 293T cells were cultured in Dulbecco’s Modified Eagle Medium (DMEM; LifeTechnologies) supplemented with 10% calf serum in a humidified 37°C incubator with 5% CO2. For the pull-down assays shown in Figure 5, cells were transfected, using X-tremeGene HP DNA Transfection Reagent (Roche), with 10 μg of pcDNA-GST, pcDNA-GST-WW, pcDNA-GST-WW-AARAA or pcDNA-p85α. Transfected cells were incubated for 24 hours and two plates of cells per treatment were harvested and flash frozen

Frozen cells were lysed by gentle pipetting in 200 μl ice cold LPD buffer (20 mM Tris–HCl (pH 7.1), 125 mM KOAc, 2 mM EDTA, 1 mM DTT, 12.5% (v/v) glycerol, 1% (v/v) Nonidet P-40, plus protease inhibitors) followed by rocking for 45 min at 4 °C. The extracts were clarified by centrifugation at 13,800 x g for 15 min. Total protein concentration was determined by Bradford assay. Binding assay samples contained 1 mg of clarified cell extract plus 20 μl of a 50% slurry of glutathione sepharose beads in a total of 200 μl LPD buffer. Samples were rocked for 1 hr at 4 °C, then pelleted at low speed in a minifuge. Pellets were washed 3X in ice cold Wash Buffer (LPD buffer in which the Nonidet P-40 was reduced to 0.1%) and then assayed by SDS-PAGE and immunoblotting.

### Peptide and antibodies

The 50-residue synthetic peptide used in Figure 3 was synthesized by GenScript USA Inc. Its sequence is:

[D-Arg]_8_-EQKLISEEDL-DNNSKWVKHWVKGGYYYYHNLETQEGGWDEPP.

This sequence is identical to that of the peptide that selectively inhibited tumor and cancer cell proliferation and viability in other studies [12, 22, 27, 28].

Protein A/G agarose beads for immunpreciptation (Figure 3) were purchased from Thermo Fisher Scientific, and the antibody was c-Myc monoclonal antibody 9E10 (Life Technologies, catalog number 132500).

The antibodies used in this study for immunoblotting (Figure 5) were (1) GST-Tag monoclonal antibody 26H1 (Cell Signaling Technologies catalog number 2624S); (2) PI3 Kinase p110α rabbit monoclonal antibody (Cell Signaling Technologies catalog number 4249); (3) PI3 Kinase p85, N-SH3, clone AB6 mouse monoclonal antibody (Sigma Aldrich catalog number 05-212).

### Statistical analysis

Statistical analysis of binding assay results was performed using Welch’s unequal variance t-test with two tails or Welch’s ANOVA test. Where necessary, p-values were adjusted for multiple hypothesis testing using Dunnett’s T3 multiple comparison test. The Graphpad Prism software package was used for statistical analysis.

For the experiment shown in Figure 7, background binding to GST was subtracted from each value, and then all measurements in that experiment were normalized by defining the binding of wild-type GST-WW to p110α as 100. This procedure complicates statistical analysis as the wild-type control (i.e. wild-type GST-WW binding to p110α) has a mean of 100 and a standard deviation of zero. Hence, we assessed significance based upon the distance between the mutant variant mean and the normalized value of 100, measured in units of the 95% confidence interval (CI) of the mean of the variant. We considered a distance of less than 1 CI as not significant (‘ns’ in Figure 7), a distance between 1 and 2 CI as significant (* in Figure 7), a distance between 2 and 3 CI as highly significant (** in Figure 7), and a distance greater than 3 CI as very highly significant (*** in Figure 7). A similar procedure was followed for the analysis of experiment shown in Figure 3E.

### Structural analysis

The predicted structure for human IQGAP1 is entry AF-P46940-F1 in the Alphafold database https://alphafold.ebi.ac.uk. Molecular graphics and analyses were performed with UCSF ChimeraX [90], developed by the Resource for Biocomputing, Visualization, and Informatics at the University of California, San Francisco, with support from National Institutes of Health R01-GM129325 and the Office of Cyber Infrastructure and Computational Biology, National Institute of Allergy and Infectious Diseases.

## Funding

This work was supported by (1) the National Cancer Institute of the National Institutes of Health under award number P30CA062203, and the UC Irvine Comprehensive Cancer Center using UCI Anti-Cancer Challenge funds; (2) the National Center for Research Resources and the National Center for Advancing Translational Sciences, National Institutes of Health, through Grant UL1 TR001414; (3) University of California Cancer Research Coordinating Committee award CTR-20-637218.

The content of this paper is solely the responsibility of the authors and does not necessarily represent the official views of the funding agencies.

## Author Contributions (CRediT format)

AJ Bardwell: conceptualization, formal analysis, investigation, methodology, supervision, validation, visualization, writing.

M Paul: investigation.

KC Yoneda: investigation.

MD Andrade-Ludeña: investigation. OT Nguyen: investigation.

DA Fruman: conceptualization, formal analysis, funding acquisition, supervision, resources.

L Bardwell: conceptualization, formal analysis, funding acquisition, investigation, methodology, project administration, supervision, validation, visualization, writing.

## Conflict of Interest Statement

The authors declare that they have no conflict of interest.

## Supplementary Figures

**Supplementary Figure 1.**
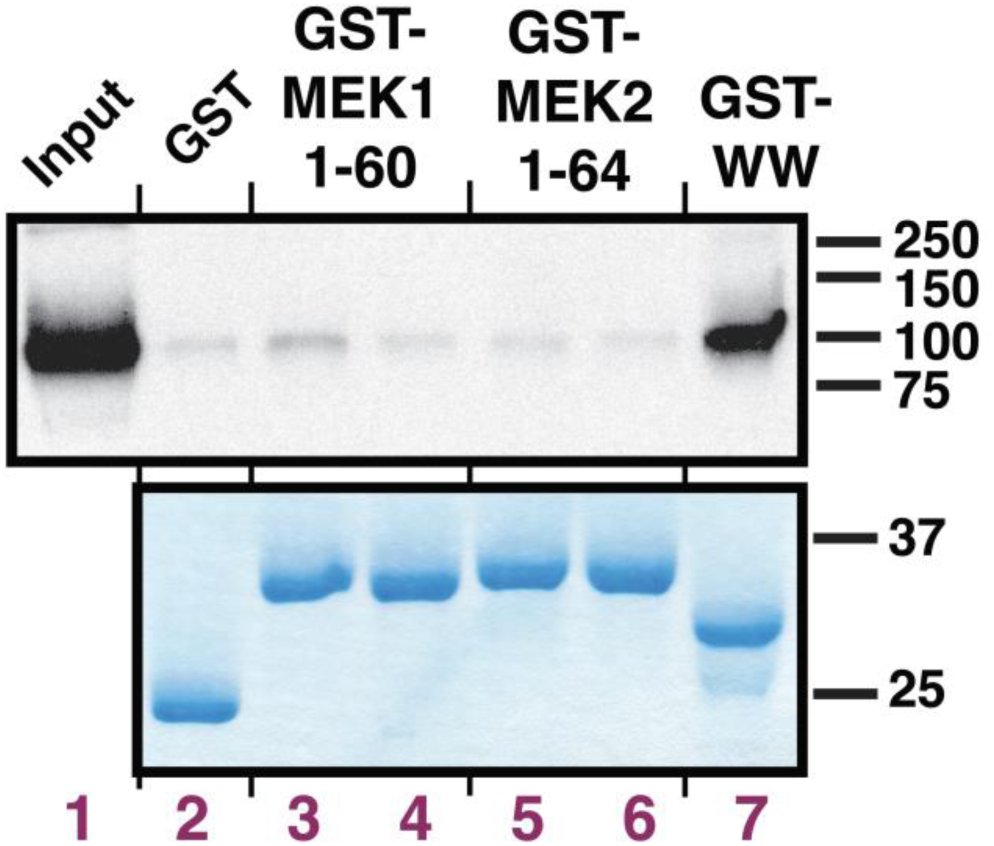
Specificity control: p110α does not bind to the N-terminal domains of MEK1 or MEK2. ^35^S-radiolabeled full-length human p110α protein was prepared by *in vitro* translation and partially purified by ammonium sulfate precipitation, and a portion (5% of the amount added in the binding reactions) was resolved on a 12% SDS-polyacrylamide (SDS-PAGE) gel (lane 1). Portions (∼1 pmol) of the same protein were incubated with 25 μg of the indicated GST fusion proteins bound to glutathione-Sepharose beads (lane 2-7), and the resulting bead-bound protein complexes were isolated by sedimentation and resolved by 12% SDS-PAGE on the same gel. The gel was analyzed by staining with a Coomassie-blue-based reagent for visualization of the bound GST fusion protein (lower panel) and by X-ray film exposure for visualization of the bound radiolabeled protein (upper panel). The migration of molecular weight markers is indicated on the right.

**Supplementary Figure 2.**
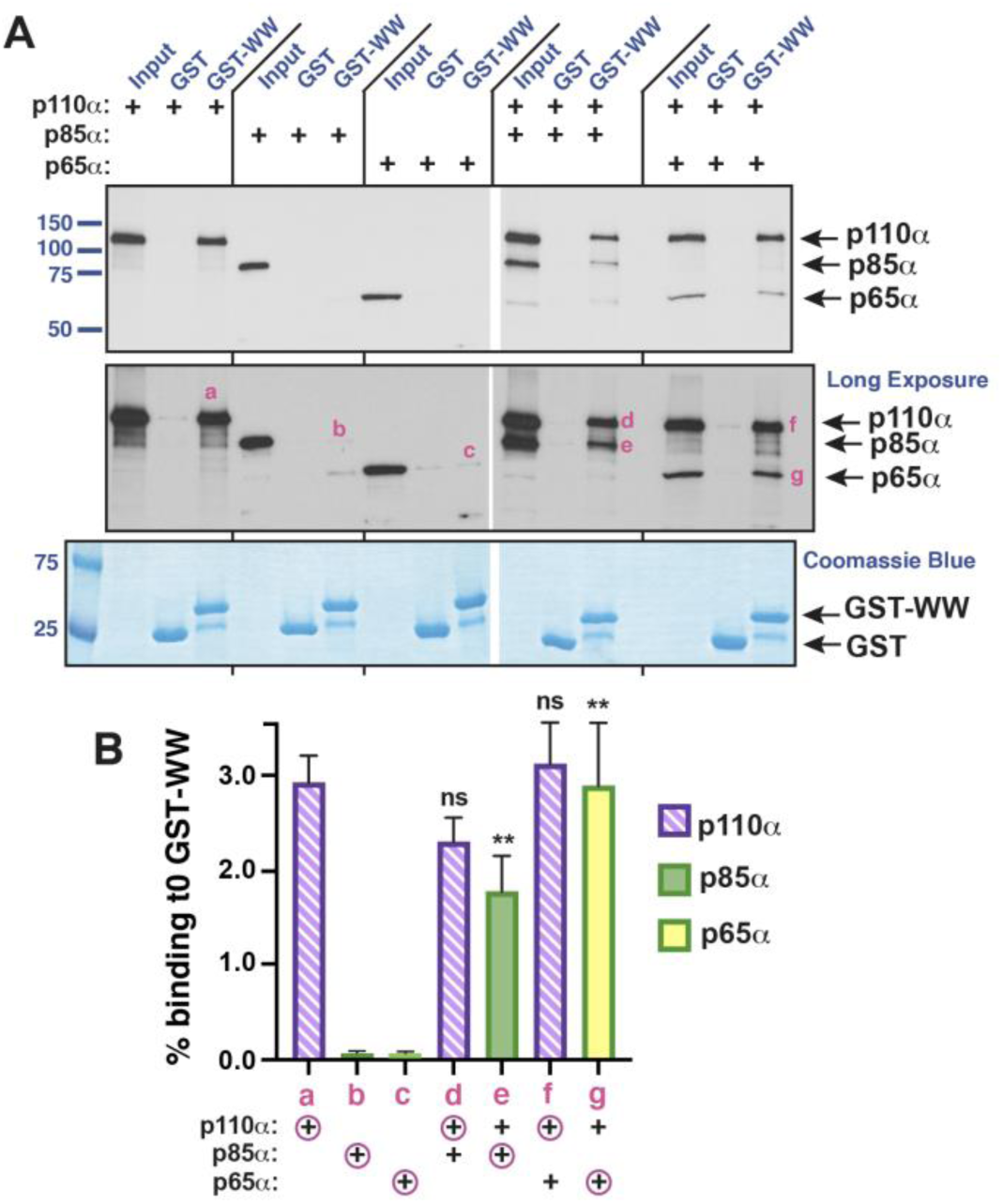
The WW domain of IQGAP binds to p110α/p85α and p110α/p65α heterodimers. (**A**) Human p110α p85α, and p65α were produced by coupled *in vitro*-transcription and translation (TNT) in separate reactions. In addition, p110α + p85α were both translated in the same reaction, and p110α + p65α were both translated in the same reaction. The products of these 5 TNT reactions were then each tested for binding to GST or GST-WW. The figure shows an autoradiogram of a representative experiment (the experiment is different from that shown in Figure 4B and 4D). Top panel, standard exposure; middle panel, longer exposure; bottom panel, Coomassie-blue-based staining to visualize GST and GST-WW. Other details as in Figure 1. The pink lower case letters a-g in the middle panel indicate bands representative of those quantified in **B**. (**B**) Quantification of the binding of human p110α, p85α and p65α to GST-WW. Data are an average of 6-10 independent repetitions of the binding assay described above and depicted in Figure 4A, measured as percent of input bound. The lowercase letters a-g below the graph match up with bands labeled in the middle panel of **A**: ‘a’ is p110α alone binding to GST-WW, ‘d’ is p110α binding to GST-WW when co-TNT’d with p85α, etc. The circled ‘+’ signs indicate which protein band is being quantified in the bar graph above. GST alone backgrounds have been subtracted from all points. Error bars show the standard error of the mean. Significance estimates are shown for the following pairwise comparisons: d to a, e to b, f to a, and g to c. **, p < 0.01; ns, not significant (p > 0.05).

**Supplementary Figure 3.**
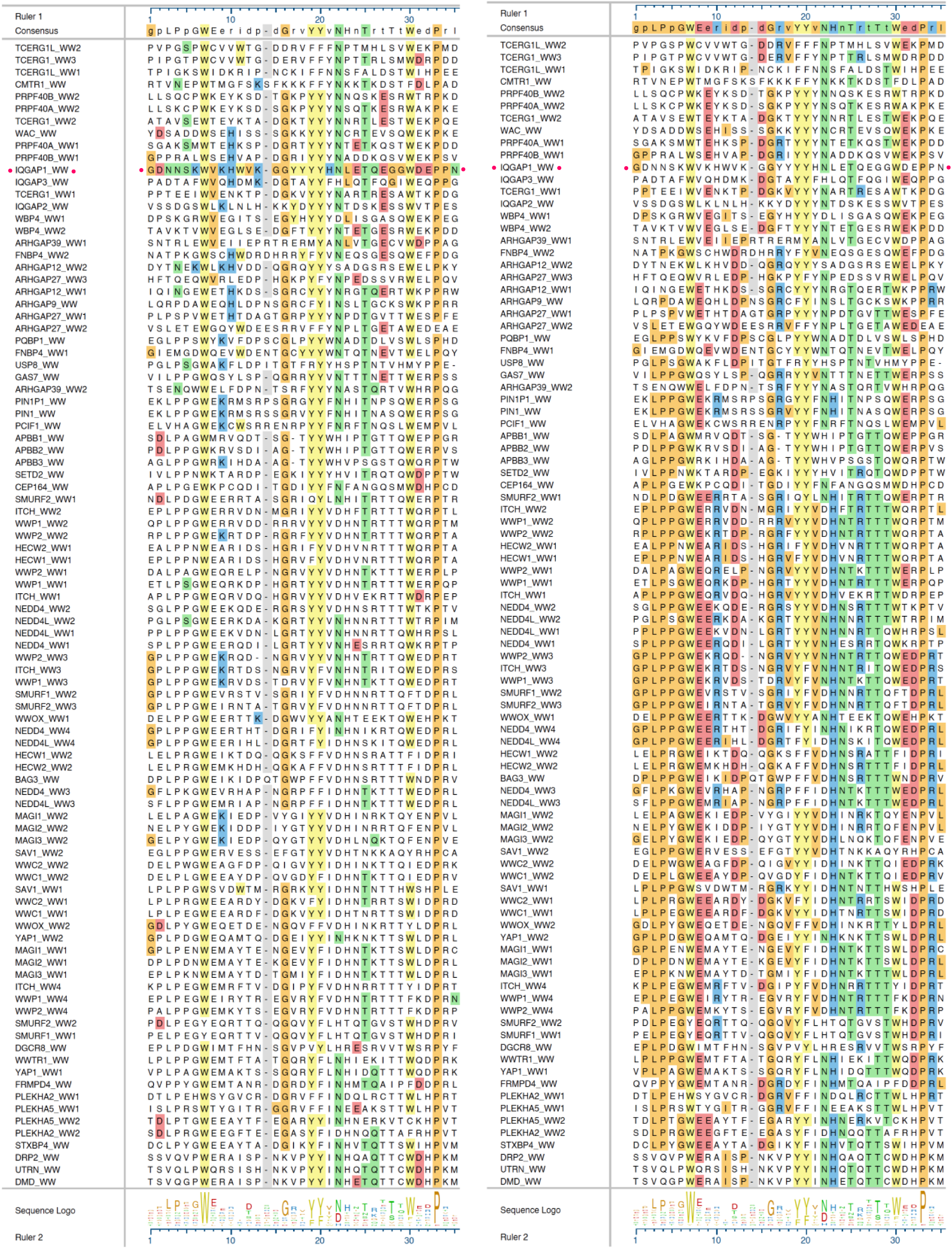
Alignment of 94 human WW domains. The sequences are listed in the order shown in the Tree in Supplementary Figure 4. The left and right alignments are identical except for coloring. On the left, residues are colored if they match IQGAP1; on the right, residues are colored if they match the consensus shown on top. The color scheme is as follows: aromatic residues (F, Y, W) are yellow, acidic residues (D, E) are red, nonpolar residues (A, G, I, L, M, P, V) are orange, polar residues (C, N, Q, S, T) are green. The numbering on the rulers are different than the numbering in Figures 6A and 7A. To convert, subtract 6 from the ruler numbers to the left of the gap in the IQGAP1 sequence and subtract 7 from the ruler numbers to the right of the gap. The alignment was generated using MegAlign Pro version 17.3.1 (DNASTAR, Inc.) using the MAFFT L-ins-I algorithm with a gap open penalty of 3 and a gap extension penalty of 0. BLOSUM62 was used as the scoring matrix.

**Supplementary Figure 4.**
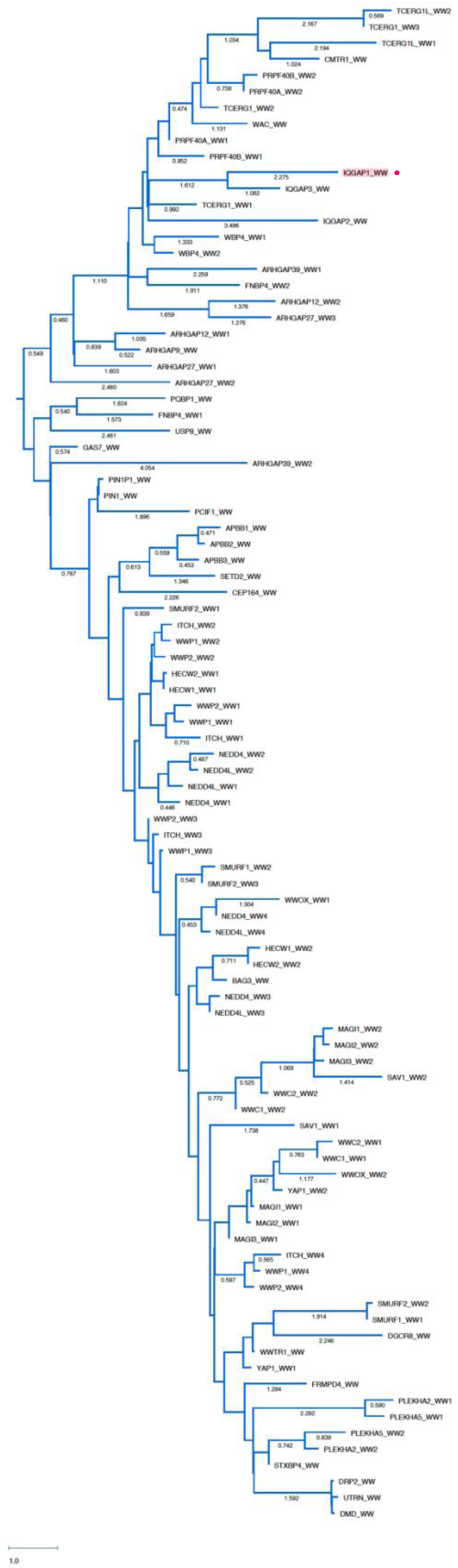
Sequence tree of 94 human WW domains. Sequence identity tree generated from the alignment shown in Supplementary Figure 3, using the maximum likelihood method with uncorrected pairwise distance and pairwise gap removal options. Other details as in Supplementary Figure 3.

**Supplementary Figure 5.**
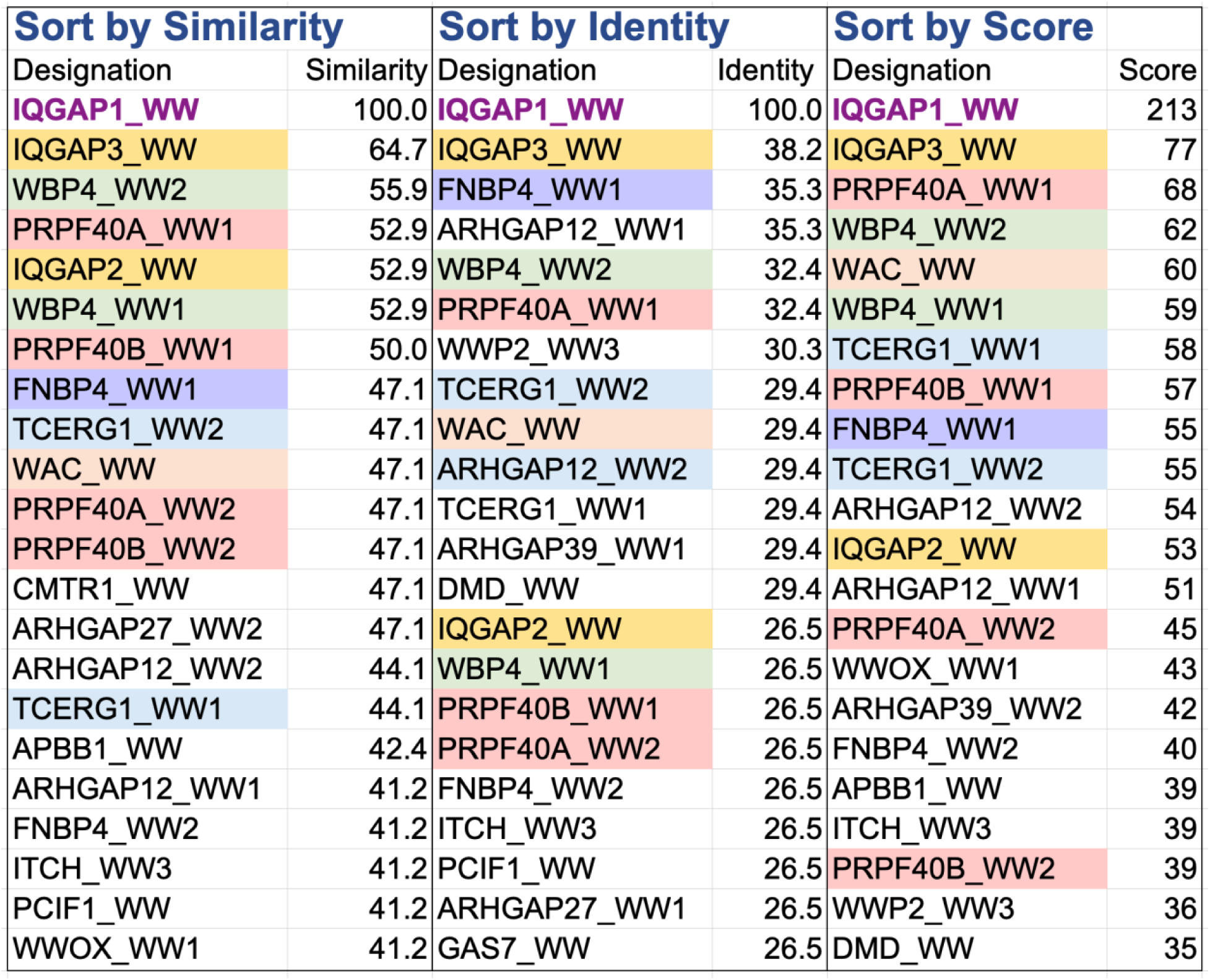
Human WW domains most closely related to the WW domain in human IQGAP1. Relatedness to IQGAP1’s WW domain was assessed by three methods: sequence similarity (left columns), sequence identity (middle columns), and the score generated by a Needleman-Wunsch global pairwise alignment with a gap penalty of 10, a gap extension penalty of 10, and BLOSUM62 used as the scoring matrix. The top 25 in each category are shown. Select WW domains are color coded as a visual aid. The sequences used are the same as shown in Supplementary Figure 3.

**Supplementary Figure 6.**
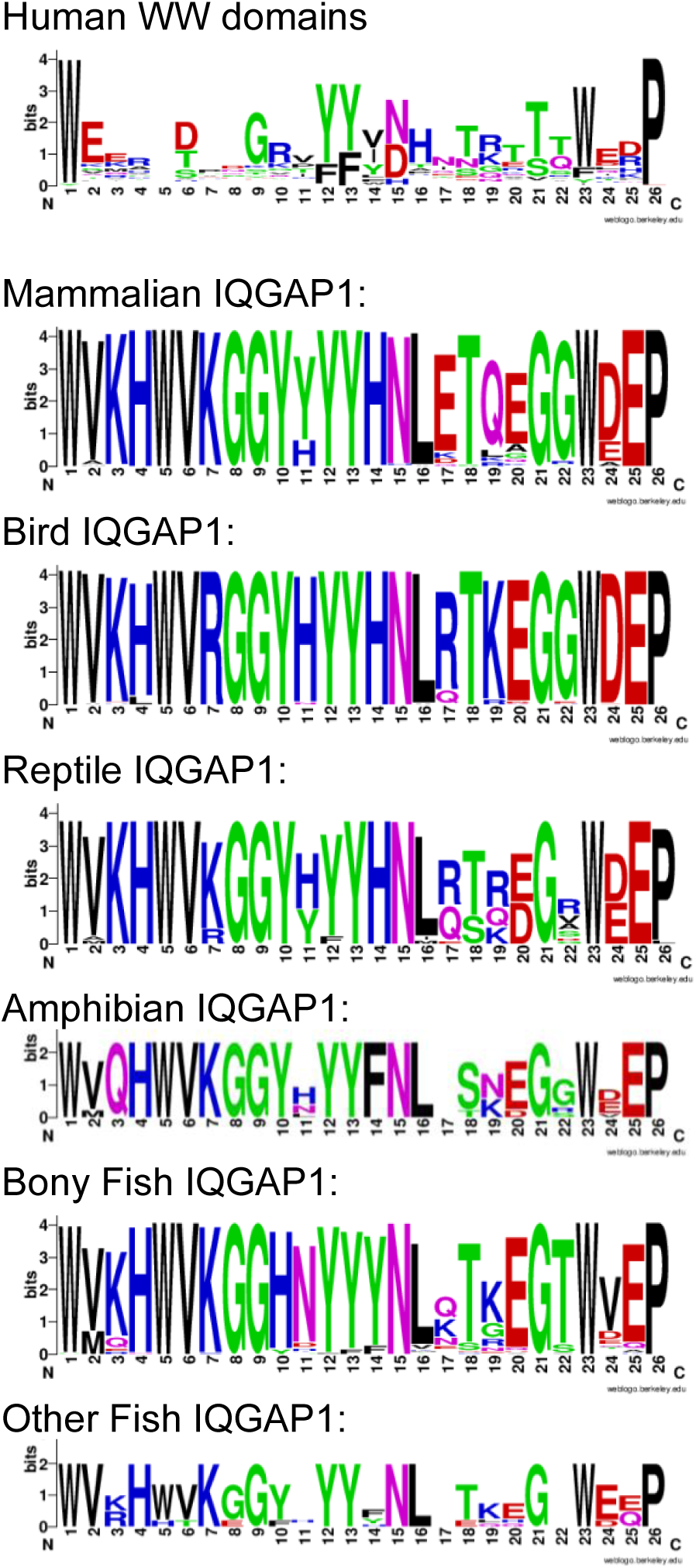
Sequence logos of human WW domains and of vertebrate IQGAP1 orthologs. The sequence logos were created with WebLogo [91]. Residue numbering as in Figures 6A and 7A. Vertebrate ortholog sequences used to create the logos are from the National Center for Biotechnology Information (NCBI) page on IQGAP1 orthologs: https://www.ncbi.nlm.nih.gov/gene/8826/ortholog/?scope=89593.

**Supplementary Table 1.** Binding assay data for WW-p110α interaction.

| Experiment <sup>a</sup> | Binding % <sup>b</sup> | $K_d$, μM <sup>c</sup> |
| --- | --- | --- |
| 1 | 5.1 | 73 |
| 2 | 5.0 | 75 |
| 3 | 4.9 | 76 |
| 4 | 4.9 | 76 |
| 5 | 4.7 | 80 |
| 6 | 4.5 | 83 |
| 7 | 4.1 | 92 |
| 8 | 3.8 | 100 |
| 9 | 3.6 | 105 |
| 10 | 3.6 | 105 |
| 11 | 3.4 | 112 |
| 12 | 3.4 | 112 |
| 13 | 3.1 | 123 |
| 14 | 2.7 | 142 |
| 15 | 2.5 | 153 |
| 16 | 2.2 | 175 |
| 17 | 2.1 | 183 |
| 18 | 2.0 | 193 |
| 19 | 1.9 | 203 |
| 20 | 1.7 | 227 |
| 21 | 1.7 | 227 |
| **Mean** | **3.4** | **129** |
| **Median** | **3.4** | **112** |
| **Standard Deviation** | **1.2** | **52** |
| **Standard Error** | **0.26** | **11** |
| **95% Confidence Interval** | **2.8-3.9** | **105-153** |

<sup>a</sup>Binding reactions (200 μl) contained ~ 1 pmole (~ 5 nM) <sup>35</sup>S-labeled, *in vitro*-translated, full-length p110α protein and 25 μg (3.9 μM) GST-WW fusion protein. The experiments are sorted by observed $K_d$ from lowest to highest.
<sup>b</sup>Percent of the input <sup>35</sup>S-labeled protein which bound to the GST fusion protein.
<sup>c</sup>Dissociation constant ( $K_d$ ), in μM, calculated based on the known input concentrations and percent binding, as described elsewhere [87, 92]. Lower $K_d$ 's indicate tighter binding.

**Supplementary Table 2.**
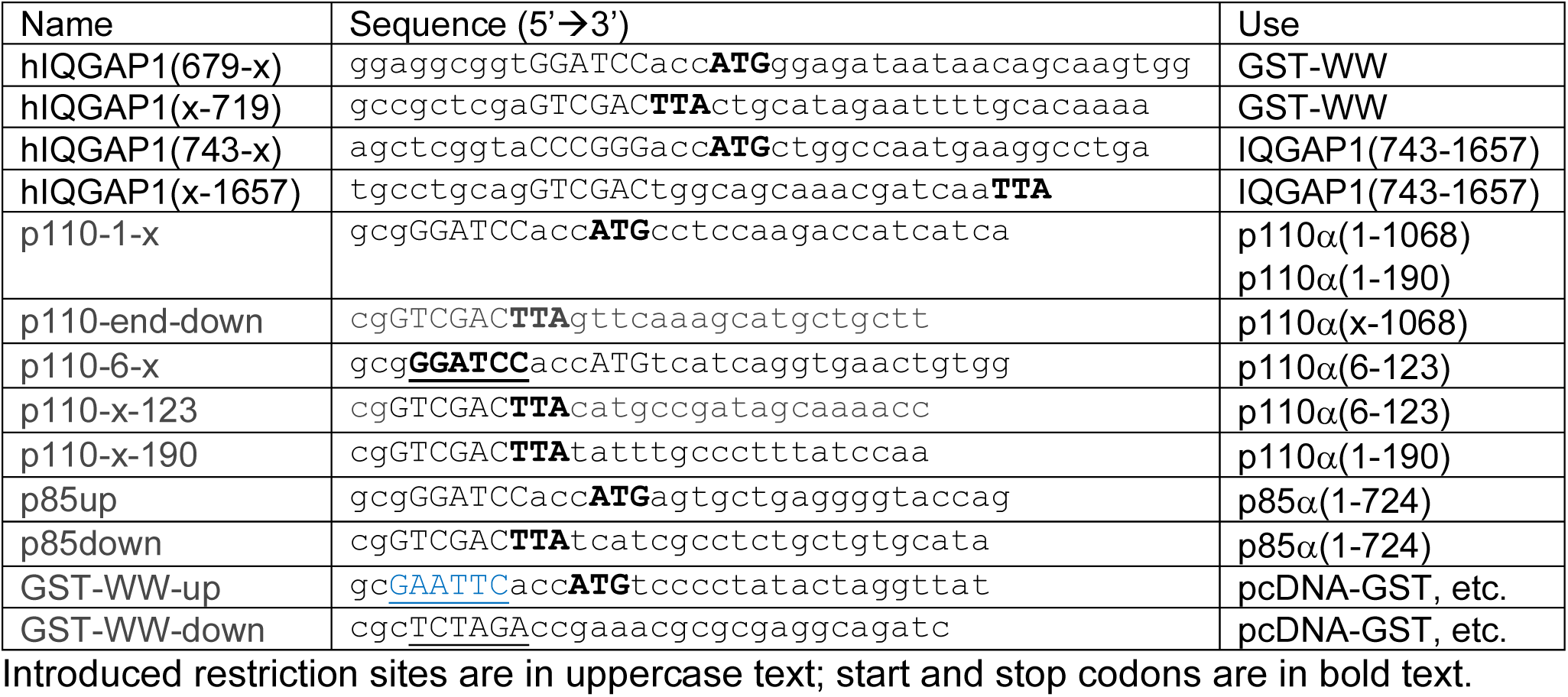
PCR primers used in this study.

## Notes

### Competing Interest Statement

The authors have declared no competing interest.

### Summary of Updates

Added 2 new figures as well as additional parts of other figures and additional data. Added new text describing the new results.

